# A riboswitch-controlled manganese exporter (Alx) tunes intracellular Mn^2+^ concentration in *E. coli* at alkaline pH

**DOI:** 10.1101/2023.05.07.539761

**Authors:** Ravish Sharma, Tatiana V. Mishanina

## Abstract

Cells use transition metal ions as structural components of biomolecules and cofactors in enzymatic reactions, making transition metals vital cellular components. The buildup of a particular metal ion in certain stress conditions becomes harmful to the organism due to the misincorporation of the excess ion into biomolecules, resulting in perturbed enzymatic activity or metal-catalyzed formation of reactive oxygen species. Organisms optimize metal concentration by regulating the expression of proteins that import and export that metal, often in a metal concentration-dependent manner. One such regulation mechanism is via riboswitches, which are 5’-untranslated regions (UTR) of an mRNA that undergo conformational changes to promote or inhibit the expression of the downstream gene, commonly in response to a ligand. The *yybP-ykoY* family of bacterial riboswitches shares a conserved aptamer domain that binds manganese (Mn^2+^). In *E. coli*, the *yybP-ykoY* riboswitch precedes and regulates the expression of two genes: *mntP*, which based on extensive genetic evidence encodes an Mn^2+^ exporter, and *alx*, which encodes a putative metal ion transporter whose cognate ligand is currently in question. Expression of *alx* is upregulated by both elevated intracellular concentrations of Mn^2+^ and alkaline pH. With metal ion measurements and gene expression studies, we demonstrate that the alkalinization of media increases cytoplasmic Mn^2+^ content, which in turn enhances *alx* expression. Alx then exports excess Mn^2+^ to prevent toxic buildup of the metal inside the cell, with the export activity maximal at alkaline pH. Using mutational and complementation experiments, we pinpoint a set of acidic residues in the predicted transmembrane segments of Alx that play a crucial role in its Mn^2+^ export. We propose that Alx-mediated Mn^2+^ export provides a primary protective layer that fine-tunes the cytoplasmic Mn^2+^ levels, especially during alkaline stress.

## Introduction

Transition metals are essential in all organisms as structural elements of proteins and RNA and as reactive centers in enzymes. Amongst these metals, Fe^2+^ acts as a cofactor in many cellular enzymes that are essential for life, e.g., those involved in respiratory pathways. During aerobic growth or in response to oxidizing agents such as hydrogen peroxide (H_2_O_2_), cells generate reactive oxygen species (ROS) that can oxidize Fe^2+^, thereby inactivating iron-dependent enzymes and leading to cytotoxic effects if not treated. To counter such ROS-caused negative consequences, *Escherichia coli* (*E. coli*) relies on isoenzymes that use Mn^2+^ instead of Fe^2+^ as a cofactor. Such enzymes protect the cell against ROS when the activity of their Fe^2+^-dependent isoenzymes is compromised (Hopkin et al., 1992). An example of such an Fe^2+^/Mn^2+^-dependent isozyme system in *E. coli* is superoxide dismutase (SOD), an enzyme that converts highly reactive superoxide radicals to molecular oxygen and H_2_O_2_: cytosolic Mn^2+^-dependent SodA takes over in aerobic conditions when the activity of Fe^2+^-dependent SodB is not sufficient to scavenge superoxide. SodC is another example of a non-iron-dependent SOD enzyme in *E. coli* that is expressed in the aerobic stationary phase and requires Cu^2+^Zn^2+^ as a cofactor to protect against ROS in the periplasmic space (Benov and Fridovich, 1994; Puget and Michelson, 1974; Strohmeier Gort et al., 1999).

To be ready for an impending ROS threat, *E. coli* maintains a constant cellular pool of Mn^2+^ (15-21 µM) through the uptake activity of its only known Mn^2+^ importer, MntH (Anjem et al., 2009; Kaur et al., 2017). MntH uses conserved acidic transmembrane residues to coordinate Mn^2+^ for import and relies on a proton gradient across the inner membrane of an *E. coli* cell as a driving force for Mn^2+^ uptake (Bozzi et al., 2019; Haemig and Brooker, 2004; Kehres et al., 2000; Makui et al., 2000). Notwithstanding its critical role within the cell, Mn^2+^ ion concentration must be limited as high concentrations of it are toxic to the cell. Excess Mn^2+^ replaces similarly sized Fe^2+^ as a cofactor in cellular enzymes and can alter levels of other metal ions (Kaur et al., 2017; Martin et al., 2015). To prevent the toxic buildup of Mn^2+^, the expression of *mntH* is repressed by elevated Mn^2+^ and an Mn^2+^-dependent transcriptional regulator MntR (Patzer and Hantke, 2001; Waters et al., 2011). As an additional layer of protection, excess Mn^2+^ is transported out of *E. coli* by its only exporter characterized to date, MntP (Martin et al., 2015; Waters et al., 2011). Similar to MntH, several conserved acidic residues within the membrane are implicated in the Mn^2+^ efflux activity of MntP (Zeinert et al., 2018). Like with *mntH*, the expression of *mntP* is regulated at the transcriptional and post-transcriptional levels by Mn^2+^ (Dambach et al., 2015).

One of the mechanisms by which *mntP* expression is tuned in response to the changing intracellular [Mn^2+^] is via the riboswitch in the 5’ untranslated region (UTR) of the *mntP* gene. Riboswitches are *cis*-acting elements in the UTRs of mRNAs, meaning that they alter transcriptional and/or translational outcomes for that mRNA. Riboswitches do so by shifting their structural ensembles upon binding to a ligand (Serganov and Nudler, 2013). For example, ligand binding might favor folding of the riboswitch RNA into a hairpin structure that terminates transcription to attenuate expression of the downstream gene (“transcriptional riboswitch”, e.g., an Mg^2+^-sensing M-box riboswitch that controls the expression of bacterial Mg^2+^ transporters *mgtA* and *mgtE* (Cromie et al., 2006; Dann et al., 2007). Alternatively, ligand binding can promote the formation of the mRNA with a single-stranded ribosome binding site (RBS), thus enhancing the translation of that mRNA (“translational riboswitch”). The *mntP* riboswitch was characterized as a translational riboswitch where the translation is turned on in response to increased intracellular [Mn^2+^] (Dambach et al., 2015). As a member of the ubiquitous riboswitch family (*yybP*-*ykoY*) (Breaker, 2022; Meyer et al., 2011), the *mntP* riboswitch regulates translation initiation on the *mntP* mRNA by binding Mn^2+^ and disfavoring formation of a stem-loop structure that sequesters the RBS of *mntP* mRNA. *In vitro* K_d_ measurements for binding to the *mntP* riboswitch vary from a low nM for aptamer-only (Bachas and Ferré-D’Amaré, 2018) to µM for a full-length riboswitch (Kalita et al., 2022). A second *yybP*-*ykoY* riboswitch in *E. coli* precedes a gene (*alx*) that, curiously, is highly induced in response to alkaline pH (Bingham et al., 1990). The expression of both *mntP* and *alx* increases in media with elevated [Mn^2+^] (Dambach et al., 2015). The *alx* encodes a putative Mn^2+^ transporter that belongs to the TerC superfamily of proteins (Anantharaman et al., 2012; Zeinert et al., 2018); however, the function of the Alx protein has not been definitively established.

A prior study indicated that overexpression of Alx results in an increase in the intracellular [Mn^2+^] and suggested that Alx may act as an Mn^2+^ importer (Zeinert et al., 2018). This proposal, however, is contradicted by the observations from earlier reports that expression of *alx* and *mntP* (Mn^2+^ exporter) are increased by elevated [Mn^2+^] in the media whereas expression of *mntH* (Mn^2+^ importer) is repressed. If Alx were indeed an Mn^2+^ importer, its expression in response to changing [Mn^2+^] would have paralleled that of MntH, not MntP. Here, we present evidence that Alx is an *exporter* of Mn^2+^ that serves as the first line of defense against the potential buildup of cytoplasmic Mn^2+^ at alkaline pH. By examining the effect of alkaline intracellular pH and elevated [Mn^2+^] on *alx* expression through transcriptional and translational reporters, we establish the link between these two environmental cues. Additionally, we demonstrate that Alx activity is stimulated by alkaline pH and posit involvement of transmembrane acidic residues of Alx in Mn^2+^ export. Our work expands the repertoire of known metal ion transporters with a transporter (Alx) that displays an alkaline pH-dependent transport activity, which may be paradigmatic of a special class of transporters responsive to multiple environmental signals.

## Results

### Both alkaline pH and increased intracellular Mn^2+^ concentration enhance *alx* expression

To study the connection between alkaline pH and Mn^2+^ homeostasis, we employed transcriptional and translational *lacZ* reporter fusions of *alx* and *mntP* cloned with their respective native promoters into single copy plasmids (Fig. 1A). Effects of extracellular alkaline pH and elevated [Mn^2+^] on gene expression were measured by β-galactosidase assays performed in *E. coli* strains that lack *alx* (referred to as Δ*alx*). The Δ*alx* strain was transformed with plasmids containing either *alx* or *mntP* transcriptional or translational reporters (Fig. 1A, Table 2). The strains containing reporter plasmids were cultivated in (i) neutral pH (LBK pH 6.8) or alkaline pH (LBK pH 8.4) media to test the effect of pH on *alx* transcriptional and translational reporters as described in prior work (Nechooshtan et al., 2009), and (ii) LB (pH 6.8) with supplemented MnCl_2_ to test the effect of Mn^2+^ on *alx* transcriptional and translational reporters. These experimental conditions were also tested on *mntP* transcriptional and translational reporters – a necessary control since the *mntP* riboswitch is only responsive to elevated [Mn^2+^]. We observed that *alx* transcription increased 5-fold at alkaline pH (Fig. 2A), whereas the *mntP* transcriptional reporter displayed a 1.5-fold induction in alkaline pH (Fig. 2A), consistent with an increase in the rate of nucleotide addition as pH increases (Mishanina et al., 2017; Stephen and Mishanina, 2022). The higher increase in the *alx* transcriptional reporter activity at alkaline pH compared to *mntP* can be explained by the proposed intrinsic terminator in hairpin D forming within the 5’UTR of *alx* in neutral but not alkaline pH (Nechooshtan et al., 2009; Stephen and Mishanina, 2022; Fig. S1). In contrast, the *alx* translational reporter produced a striking 68-fold higher signal in alkaline pH, whereas the *mntP* translational reporter was unaffected by alkaline pH (Fig. 2B). These results indicate that *alx* expression is largely regulated post-transcriptionally in alkaline pH, consistent with previous work (Nechooshtan et al., 2009).

**Figure 1.**
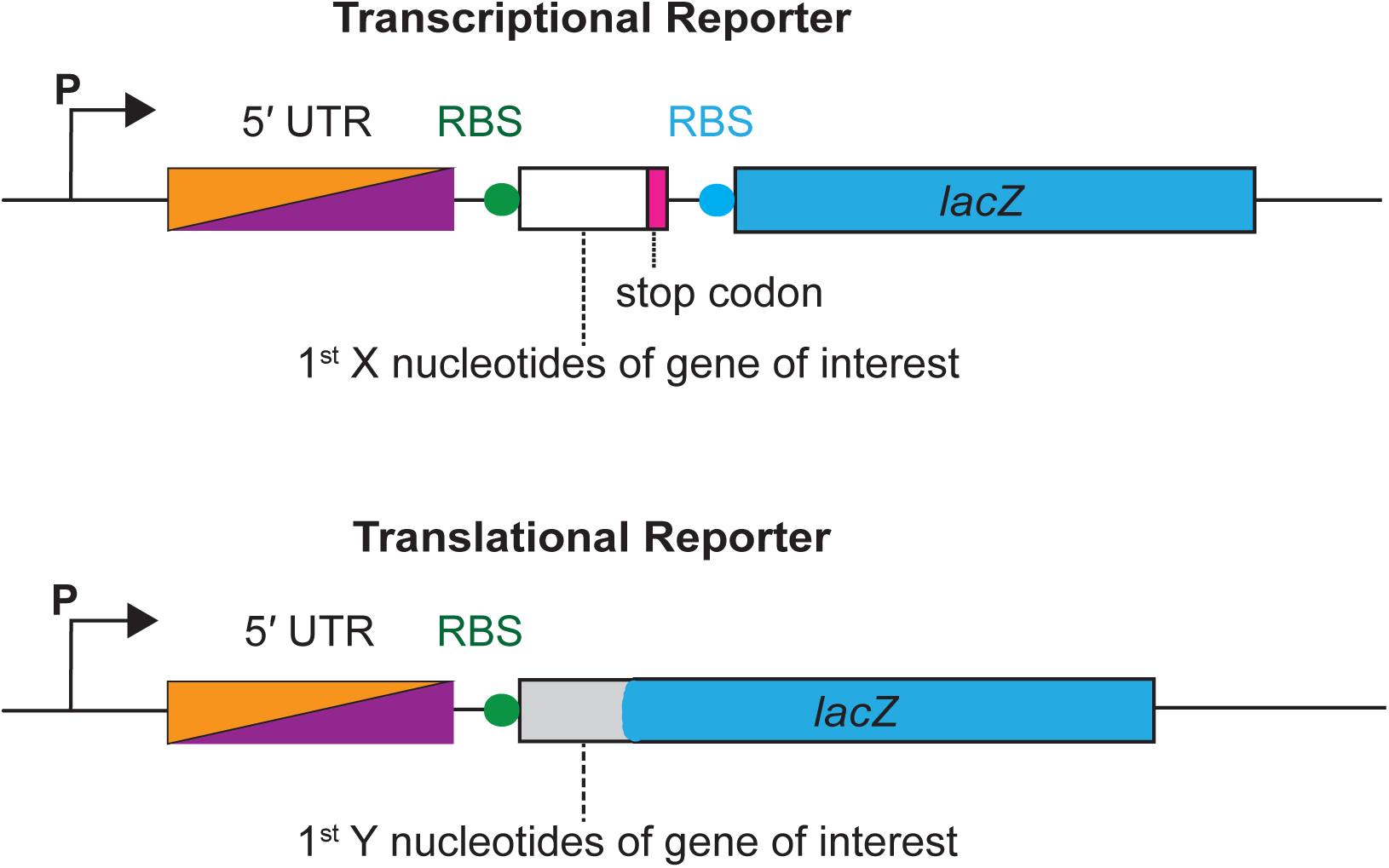
A schematic illustration of transcriptional and translational reporters of *alx* and *mntP.* A transcriptional reporter contains a promoter (P) of *alx* or *mntP,* followed by the 5’ UTR (PRE in the case of *alx)*, a ribosome binding site (RBS) of the gene of interest indicated by a filled circle in green, the first X nucleotides of the gene of interest (53 and 47 nt for *alx* and *mntP,* respectively), and a stop codon indicated by a rectangular box in magenta, followed by *lacZ* gene with its own RBS indicated by a rectangular box and filled circle respectively in blue. A translational reporter contains a promoter (P) of *alx* or *mntP* is followed by a 5’ UTR element (PRE in the case of *alx*), an RBS of the gene of interest indicated by a filled circle in blue, and first Y nucleotides of the gene of interest (99 and 47 nt for *alx* and *mntP*, respectively) fused to in frame with the 8th codon of *lacZ* gene.

**Figure 2.**
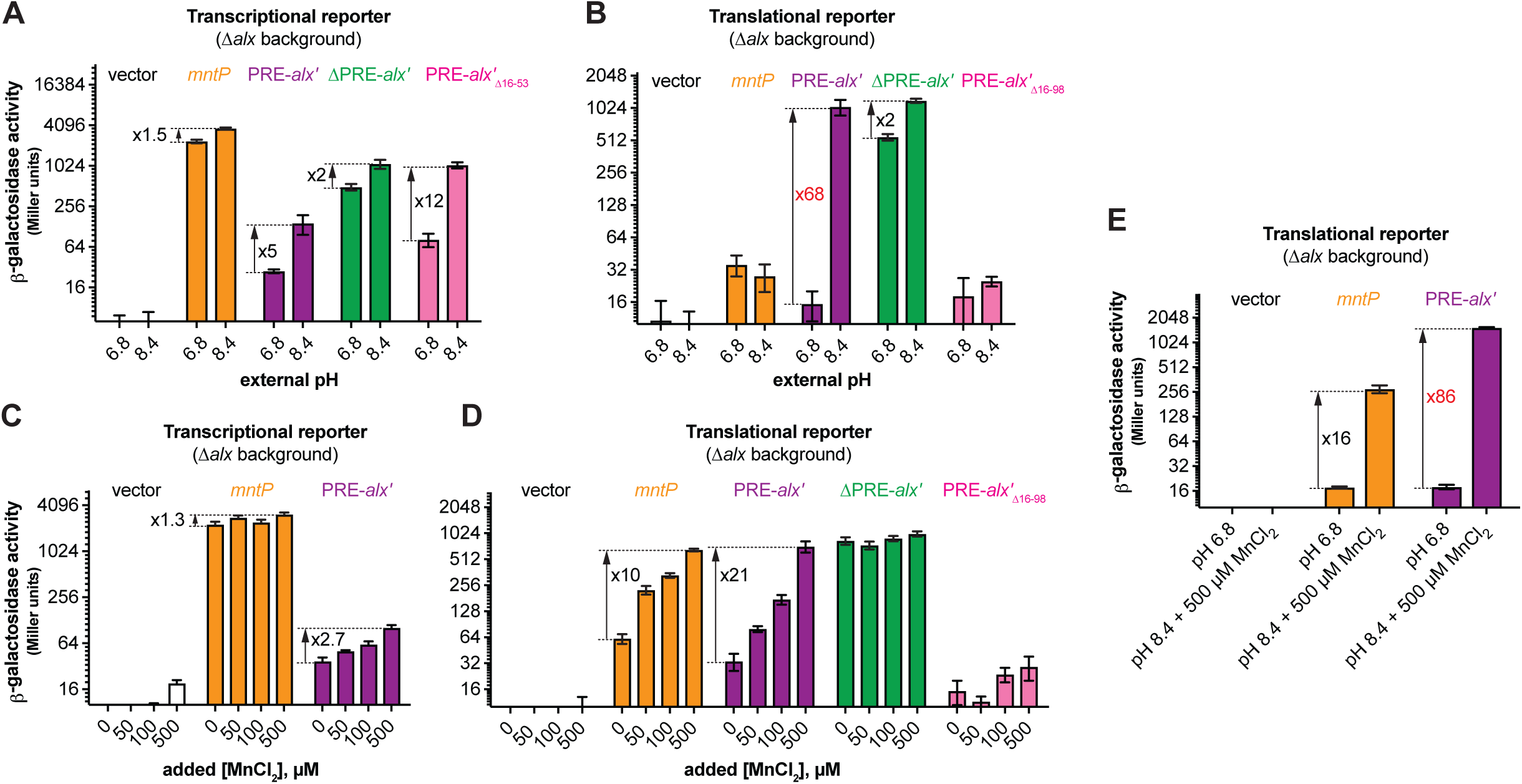
Regulation of *alx* expression by PRE in response to alkaline pH and elevated [Mn^2+^]. Shown in panels A and C are the β-galactosidase activities (Miller units) of mid-log phase grown cultures of Δ*alx*::Kan derivatives of MC4100 strain of *E. coli* (RAS31) carrying one of the following plasmids: promoter-less vector with *lacZ* (pMU2385), transcriptional reporter of *mntP* (*P_mntP_-* 5’UTR*-mntP’*-*`lacZ`*, PRA48), transcriptional reporter of *alx* (P*_alx_*-PRE*-alx’-lacZ*, PRA40), ΔPRE derivative of *alx* transcriptional reporter (P*_alx_*-alx’-*lacZ,* pRA41), or a transcriptional reporter of *alx* that lacks putative transcriptional pause sites in the vicinity of the first *alx* codon (alx’_Δ16-53_, PRA49). Above cultures were cultivated in LBK media with pH 6.8 and pH 8.4 (panel A) and pH 6.8 LB with and without supplemented MnCl_2_ (panel C). Similarly, shown in panels B and D are the β-galactosidase activities of mid-log phase grown cultures of *Δalx*::Kan derivatives of MC4100 strain of *E. coli* (RAS31) carrying `one` of the following plasmids: promoter-less vector with *lacZ* (pMU2386), translational reporter of *mntP* (P*_mntP_*-5’UTR-*mntP’-lacZ,* PRA57), translational reporter of *alx* (P*_alx_*-PRE-*alx’-lacZ*, PRA54), ΔPRE derivative of *alx* translational reporter (P*_alx_*-alx’-*lacZ,* PRA55), or translational reporter of *alx* that lacks putative transcriptional pause sites in the vicinity of the first *alx* codon (alx’_Δ16-98_, PRA56). Above cultures were cultivated in LBK media with pH 6.8 and 8.4 (panel B) and pH 6.8 LB with and without supplemented MnCl2 (panel D). A combined effect of alkaline pH and supplemented MnCl_2_ on translational reporters is illustrated in panel E. The error shown is standard deviation of three repeats of the experiment.

The 5’ UTR of *alx* mRNA referred to as the pH-responsive RNA element (PRE) regulates *alx* translation in response to a pH change (Nechooshtan et al., 2009. We observed that the translational reporter of *alx* that lacks PRE (ΔPRE) exhibits only a 2-fold increase in alkaline pH vs 68-fold increase with PRE present (Fig. 2B). PRE contains two intrinsic transcription terminators (Fig. S1), and their absence is the dominant cause of higher transcription output in the ΔPRE transcriptional reporter. In addition to riboswitch regulation, bacterial translational output also appears to be tuned by the presence of transcriptional pause sites. Specifically, the analysis of cellular nascent elongating transcripts (NET-seq) isolated from immunoprecipitated RNA polymerases (RNAPs) revealed that transcriptional pause sites are generally enriched in the vicinity of the translation start sites in *E. coli* and *B. subtilis* (Larson et al 2014). We observed seven prominent transcriptional pauses on analysis of reported NET-seq data for *E. coli* (Larson et al., 2014) in the vicinity of the translation start site of *alx* (Fig. S2). Earlier studies that employed translational reporters of *alx* lacked the last but dominant transcriptional pause sequence. The translational reporter of *alx* used in this study includes all putative transcriptional pause sequences mentioned in Fig. S2. We observed a much higher fold-change in translational reporter activity (68-fold) in comparison to the translational reporter (7.8-fold) employed in previous report (Nechooshtan et al., 2009), stressing the importance of the translation start-site proximal RNAP pauses for efficient translation initiation. We noted a 12-fold increase for the transcriptional reporter that lacks nucleotides 16-53 of *alx*’ (*alx*’_Δ16-53_) containing putative transcriptional pause sequences (Fig. S2). Absence of PRE or putative transcriptional pause sequences in *alx*’ increased transcriptional output in both pH conditions (Fig. 2A), suggesting that these sequences are important to slow down *alx* transcription. In comparison, ΔPRE *alx* translational reporter displayed high β-galactosidase activity in both pH conditions, with a 2-fold increase in β-galactosidase activity in alkaline vs neutral pH (Fig. 2B), in good agreement with the corresponding 2-fold increase in ΔPRE *alx* transcription upon alkalinization (Fig. 2A). Taken together, these results suggest that PRE regulates both transcription and translation of *alx* in a pH-responsive manner. Surprisingly, there was very low reporter activity in case of translational reporter fusion of *alx* bearing *alx*’_Δ16-98_, suggesting that the putative transcriptional pauses proximal to the *alx* translation start site are critical for *alx* translation initiation.

Upon supplementation of 500 µM MnCl_2_ in the LB media (pH 7.2), the *alx* and *mntP* transcriptional reporter outputs increased only 2.7-fold and 1.4-fold, respectively (Fig. 2C). In stark contrast, both *alx* and *mntP* translational reporters were progressively induced by increasing [Mn^2+^] in the media, with a 21-fold and 10-fold increase in the reporter activity, respectively, at the highest MnCl_2_ concentration tested (500 µM) (Fig. 2D). These results suggest that expression of both *alx* and *mntP* is post-transcriptionally responsive to elevated extracellular [Mn^2+^]. The ΔPRE translational reporter of *alx* was unaffected by the supplemented MnCl_2_ and displayed high reporter activity throughout (Fig. 2D), likely due to the absence of premature transcription termination within PRE (Fig. S1). Collectively, these observations suggest that PRE tunes *alx* expression in an Mn^2+^ concentration-responsive manner. Similar to the pH response experiments above, the translational reporter fusion of *alx* bearing *alx*’_Δ16-98_ displayed very low β-galactosidase activity, reinforcing the notion that translation start-site proximal RNAP pauses are important for successful translation of *alx*.

We were intrigued to test the combined effect of the two environmental signals, alkaline pH and elevated [Mn^2+^], on *alx* and *mntP* translational reporter fusions. We found that *alx* and *mntP* translational fusions are induced 86-fold and 16-fold, respectively, in alkaline media supplemented with 500 µM MnCl_2_ (Fig. 2E). Altogether, our results (Fig. 2B, 2D, 2E) demonstrate that the effects of elevated extracellular [Mn^2+^] and alkaline pH on *alx* expression are additive in nature and may have independent routes to enhance *alx* expression. On the other hand, *mntP* expression is induced further if both alkaline pH and extra MnCl_2_ are provided compared to MnCl_2_ supplementation alone (Fig. 2E), even though alkaline pH had no significant impact on *mntP* expression (Figs. 2A and B). These observations indicate that alkaline pH enhances the effect of elevated [Mn^2+^] on *mntP* expression, consistent with previously published work (Kalita et al., 2022).

### Alx does not participate in maintaining cellular pH or redox homeostasis

Our study confirms previously published work which demonstrated that the expression of *alx* is induced in media with alkaline pH and elevated [Mn^2+^] (Fig. 2, (Bingham et al., 1990; Dambach et al., 2015)). Perhaps the simplest explanation for the pH stress-induced production of Alx would be its direct involvement in bringing the high intracellular pH back into its neutral physiological range. To probe the contribution of Alx to pH homeostasis, we tested the growth of the parent strain (MC4100) and its Δ*alx* derivative (RAS31) in LBK media at pH 6.8 or 8.4. The growth of the Δ*mntP* mutant and the Δ*alx* Δ*mntP* double mutant was also tested under these same conditions. In general, the growth of strains slowed in the pH 8.4 media compared to pH 6.8 (Fig. 3A). Absence of Alx did not affect the growth rate in alkaline pH in comparison to its parent strain, suggesting that Alx does not provide a growth advantage in alkaline pH.

**Figure 3.**
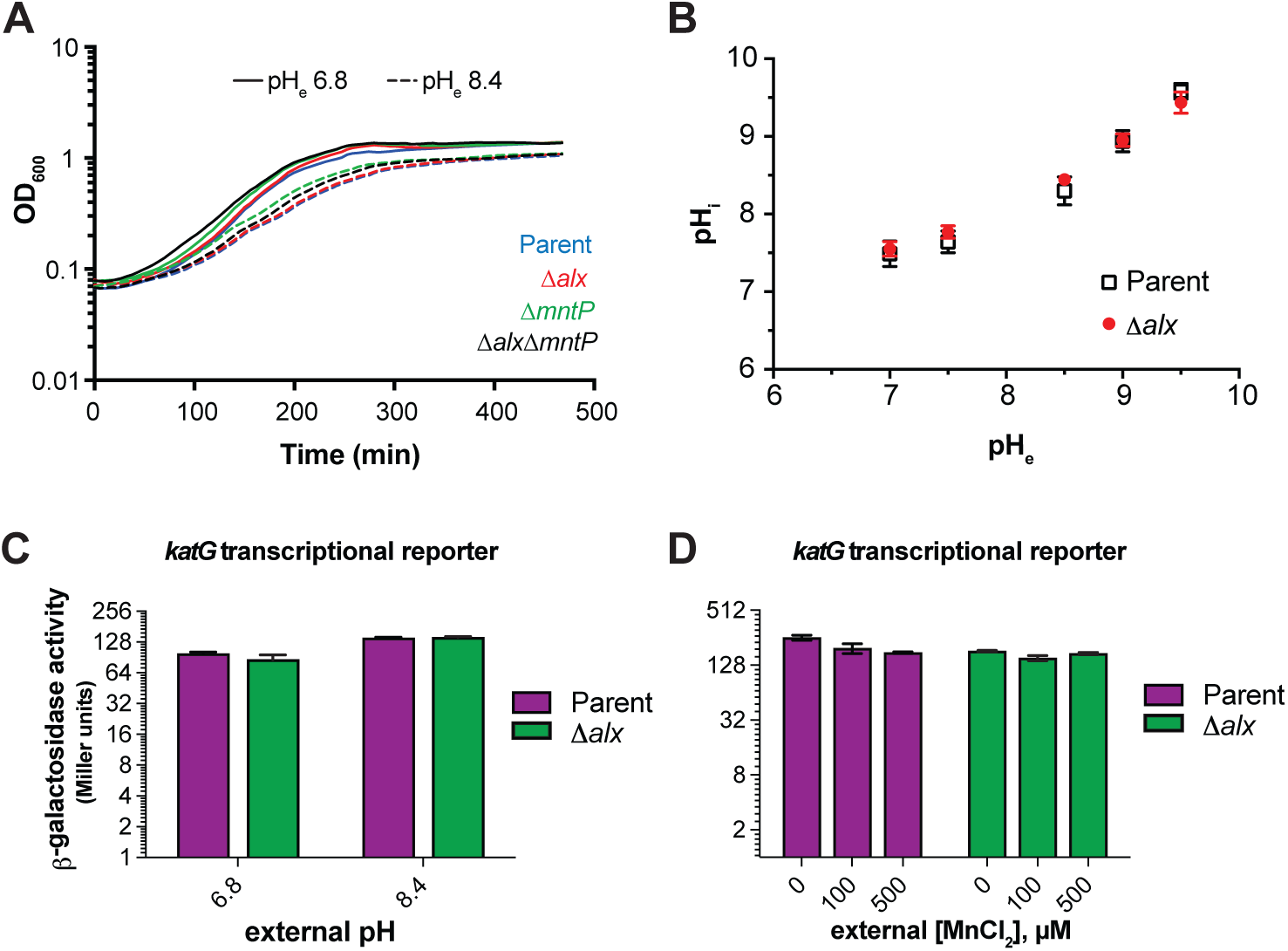
Effect of increased external pH on growth, cytoplasmic pH, and cellular oxidative stress. (A) Growth of the parent strain (MC4100) and its derivatives (*Δalx*::Kan derivative, RAS31; *ΔmntP*::Kan derivative, RAS32, and *Δalx ΔmntP*::Kan derivative, RAS42) in LBK media with pH 6.8 and pH 8.4. (B) Measurements of cytoplasmic pH in the parent strain (MC4100) and its *Δalx*:: Kan derivative (RAS31) expressing pHluorin in M63A media with varying pH. β-galactosidase activity (Miller units) as a reporter of *katG* transcription was measured for mid-log phase grown cultures of the parent strain (AL441) and its *Δalx*::Kan derivative (RAS136) cultivated in LBK media with pH 6.8 and 8.4 (panel C) or pH 6.8 LB with and without supplemented MnCl_2_ (panel D). The error shown is standard deviation of three repeats of the experiment.

To further rule out the involvement of Alx in pH homeostasis, cytoplasmic pH of the parent strain and its Δ*alx* derivative were measured in M63A media over a range of pH values (7, 7.5, 8.5, 9, and 9.5) using genetically encoded ratiometric pHluorin as a reporter (Martinez et al., 2012). Cytoplasmic pH of the parent strain increased as external pH was elevated (Fig. 3B). Cytoplasmic pH of the Δ*alx* mutant did not display any significant difference compared to its parent strain when external pH was elevated (Fig. 3B). These results indicate that Alx participation in countering alkalization of cytoplasmic pH is unlikely.

Mn^2+^ protects cells from oxidative damage by serving as a cofactor in enzymes where it can replace oxidized iron in the catalytic center to rescue activity or as a cofactor in enzymes that prevent the buildup of ROS, such as SodA. Indeed, the expression of *mntH* (Mn^2+^ importer) increased in the presence of high extracellular or endogenously produced H_2_O_2_ (Anjem et al., 2009; Kehres et al., 2002, 2000). MntH becomes vital for growth in aerobic conditions of a strain (Δ*katG* Δ*katE* Δ*ahpCF*) lacking catalase and peroxidases that would normally clear accumulating H_2_O_2_ (Anjem et al). Previously, it was noted that *alx* expression is sensitive to the presence of an oxidizing agent, paraquat (Pomposiello et al., 2001). Considering that *alx* expression is induced by alkaline pH, we investigated the connection between alkaline pH and oxidative stress. We assessed the oxidative stress in the parent strain and its Δ*alx* mutant in alkaline pH and in the presence of elevated extracellular [Mn^2+^], using *katG* (bifunctional catalase-peroxidase) transcriptional reporter, which is induced in the oxidative environment (Li and Imlay, 2018). We measured the *katG* transcriptional reporter activity in LBK media with pH 6.8 or 8.4 (Fig. 3C). Low induction (1.5-fold) of the *katG* transcription reporter was observed in both parent and its Δ*alx* derivative at pH 8.4. The Δ*alx* mutant displayed similar *katG* transcriptional reporter activity to its parent strain at both pHs. These results indicate mild oxidative stress at alkaline pH. The *katG* transcriptional reporter activity was somewhat repressed in both parent strain and Δ*alx* mutant in LB (pH 6.8) media upon supplementation of MnCl_2_ (Fig. 3D), consistent with the role of Mn^2+^ in alleviating oxidative stress. The difference in the induction of the *katG* transcriptional reporter activity in the parent vs its Δ*alx* mutant in LB media lessened with increasing Mn^2+^ concentration in the media. The mild repression of the *katG* transcriptional reporter activity in the Δ*alx* mutant compared to its parent strain will be elaborated on in the Discussion. Overall, these results suggest that Alx may not be directly participating in the maintenance of a redox state at alkaline pH or elevated extracellular [Mn^2+^].

### Alx mediates the export of Mn^2+^ in alkaline environment

A previous study reported that *mntP* encodes an exporter of Mn^2+^ and its absence makes *E. coli* growth extremely sensitive to elevated [Mn^2+^] in the media (Waters et al., 2011). We observed that the growth of the Δ*mntP* mutant and the Δ*alx* Δ*mntP* double mutant slowed in LB media (pH 6.8) supplemented with 500 µM MnCl_2_, as expected (Fig. S3). In contrast, the absence of Alx alone did not lead to any growth retardation of the strains in media with elevated [Mn^2+^] (Fig. S3), suggesting Alx does not contribute significantly to Mn^2+^ homeostasis. In testing the combined effect of pH 8.4 and 500 µM MnCl_2_ supplementation on *E. coli* growth, we observed that the growth of the Δ*mntP* mutant and the Δ*alx* Δ*mntP* double mutant slowed compared to the parent strain, whereas its Δ*alx* derivative’s growth rate was indistinguishable from the parent (Fig. S3).

Expression of both *alx* and *mntP* is controlled by the riboswitches within their 5’ UTRs, which belong to the *yybP*-*ykoY* family of transition metal ion-binding riboswitches (Barrick et al., 2004, (Dambach et al., 2015) . Like MntP, Alx is predicted to encode an inner membrane transition metal ion transporter (Daley et al). In light of these similarities and the observed upregulation of both *alx* and *mntP* by elevated [Mn^2+^] (Fig. 2D, (Dambach et al., 2015)), we were curious to test whether heterologous expression of *alx* would rescue the Mn^2+^ sensitivity phenotype of the Δ*mntP* mutant (RAS32 strain). We found that the expression of *alx* from a *trc* promoter (P*_trc_*) indeed partially rescued the growth of Δ*mntP* mutant in the presence of supplemental Mn^2+^ (Fig. 4A), whereas growth of the parent strain was not altered by the expression of *alx* from P*_trc_*. These results indicate that Alx may mediate the export of Mn^2+^ in circumstances when cytoplasmic Mn^2+^ levels are elevated. To test whether the Mn^2+^ content of the cells increases at alkaline pH when Alx is most expressed, intracellular concentrations of transition metals ions (Mn^2+^, total iron and Zn^2+^) were measured in the Δ*alx* mutant (to preclude any transport of metal ions by the Alx prior to the measurement) under neutral and alkaline pH (Fig. 4B). Overall, our measured metal ion concentrations agree with previously published values (Anjem et al., 2009; Kaur et al., 2014). An increase of 1.5-fold in the total intracellular Mn^2+^ (from 28 to 42 μM) in the Δ*alx* mutant was indeed noted at pH 8.4 when compared with pH 6.8 (Fig 4B). Importantly, we observed that although P*_trc_*-driven expression of Alx did not change total intracellular Mn^2+^ levels in the Δ*alx* mutant at pH 6.8, it did reduce the total intracellular Mn^2+^ 2-fold at pH 8.4 (from 42 to 19 μM). These results suggest that the Alx function may be to prevent the buildup of intracellular Mn^2+^ specifically under alkaline pH. In the case of the total iron content (Fe^2+^ and Fe^3+^), we noted an increase of 2-fold in the total intracellular iron of the Δ*alx* mutant at pH 8.4 vs 6.8. There was no significant change in total intracellular Zn^2+^ in the Δ*alx* mutant at pH 8.4 vs. 6.8. The P*_trc_*-driven expression of Alx did not produce a significant change in total intracellular iron or Zn^2+^ in the Δ*alx* mutant at either pH, suggesting that Alx likely has no role in the transport of iron or Zn^2+^ and is selective for Mn^2+^.

**Figure 4.**
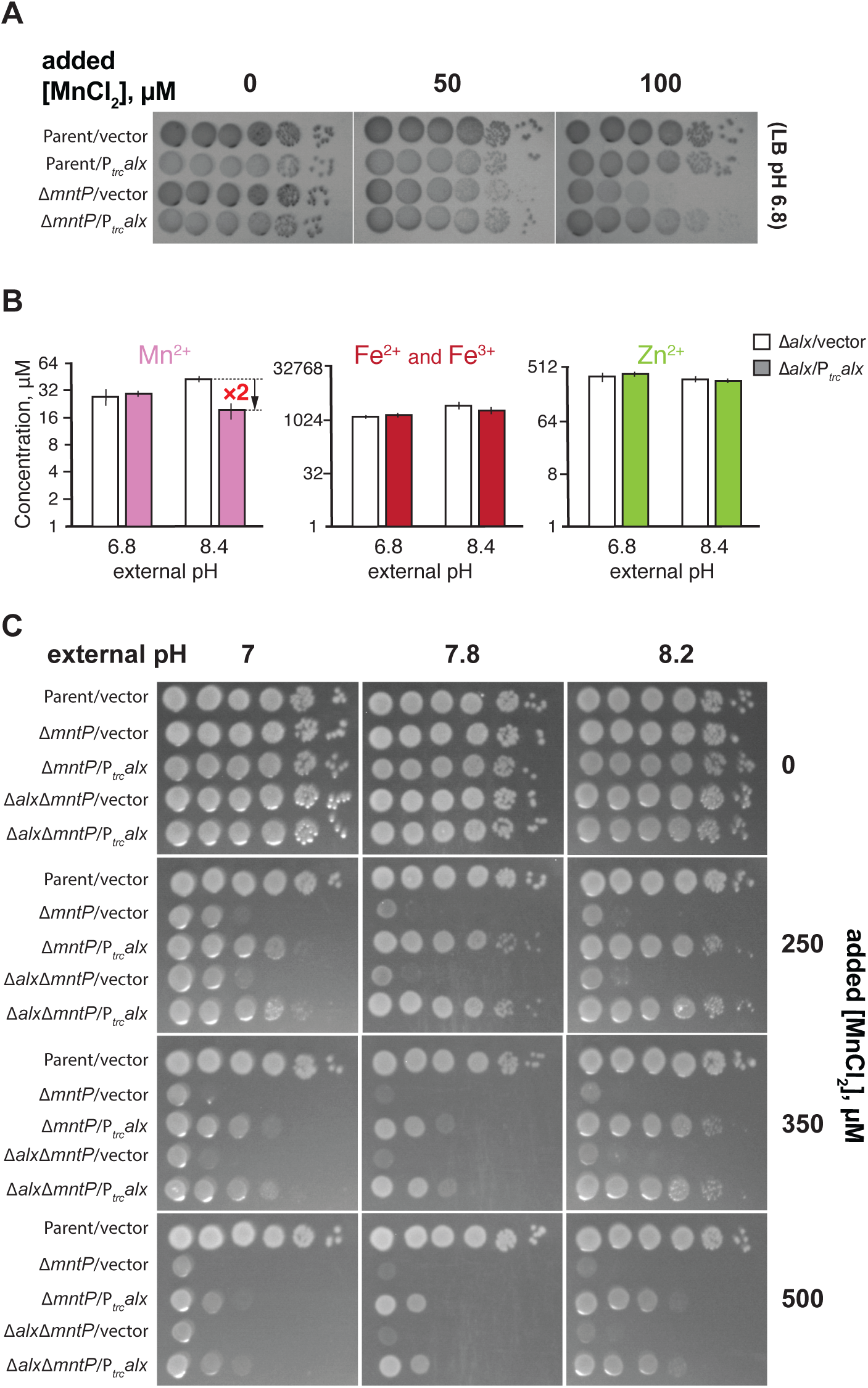
Alx exports Mn^2+^ in alkaline pH. (A) Tenfold serial dilutions of overnight-grown cultures of parent strain (MC4100) `and` its *ΔmntP*::Kan derivative, `each` bearing `an` empty vector (pHYD5001) or the same vector expressing Alx from a P*_trc_* promoter were spotted on the surface of LB agar containing the appropriate concentration of ampicillin, MnCl_2_ and IPTG. (B) Intracellular Mn^2+^, Fe, and Zn^2+^ concentrations were measured in cultures of *Δalx* mutant (RAS31) containing a vector (pHYD5001) or a derivative of pHYD5001 expressing Alx from a P*_trc_* promoter (PRA27), grown to mid-log in LBK media pHs 6.8 and 8.4 supplemented with 1 mM IPTG and ampicillin. The error shown is standard deviation of three repeats of the experiment. (C) Tenfold serial dilutions of overnight-grown cultures of parent strain (MC4100) and its derivatives (*ΔmntP*::Kan derivative, RAS32, and *Δalx ΔmntP*::Kan derivative, RAS42), each bearing an empty vector (pHYD5001) or the same vector expressing Alx from a P*_trc_* promoter were spotted on the surface of LB agar of varying pH and supplemented with appropriate concentration of ampicillin, MnCl_2_, and IPTG.

The above data indicated that P*_trc_*-driven expression of Alx decreases intracellular [Mn^2+^] in the Δ*alx* mutant at pH 8.4, but not pH 6.8, suggesting that Mn^2+^ transport by Alx is pH-dependent (Fig. 3S). To test the possibility that the mechanism of Mn^2+^ export by Alx is proton dependent, we performed assays for detecting substrate-induced proton release in inside-out vesicles using published procedures (Dubey et al., 2021). Everted membrane vesicles were prepared for the strain that contains an in-frame deletion of both *alx* and *mntP* on the chromosome (RAS42). In place of the chromosomally encoded proteins, a human influenza hemagglutinin (HA)-tagged derivative of Alx or MntP (Alx^HA^ or MntP^HA^) was expressed in the strain RAS42, from pRA50 or pRA70 plasmid, respectively. Successful expression of each tagged protein was confirmed by anti-HA Western blotting. A pH gradient across the vesicle membrane was generated via F_0_F_1_ ATPase activity by the addition of ATP to the vesicle suspension (Fig. S4). To monitor the generation of pH gradient, a pH gradient-sensitive, fluorescent dye 9-amino-6-chloro-2-methoxyacridine (ACMA) was employed. We noted an expected quenching of fluorescence upon the addition of ATP due to the generation of a proton gradient across the membrane suggesting vesicles were active. If Mn^2+^ transport by Alx or MntP is dependent on proton release, then a dequenching of ACMA fluorescence upon the addition of MnCl_2_ would be expected. However, we did not observe a significant change in the fluorescence intensity of ACMA upon the addition of MnCl_2_, suggesting that the transport of Mn^2+^ by Alx and MntP is unlikely to be accompanied by an H^+^ antiport.

We observed above that Alx expression from a *trc* promoter decreased the intracellular [Mn^2+^] in alkaline pH (Fig. 4B) without an obvious accompanying H^+^ transport (Fig. S4). Based on these findings, we speculated that alkaline pH may stimulate the Mn^2+^ export activity of Alx directly, perhaps by altering the protonation state of key Alx residues (see “A set of acidic residues in the transmembrane helices are crucial for Alx-mediated Mn^2+^ export” section of the Results below). To test this hypothesis, we probed the combined effect of elevated pH and extracellular [Mn^2+^] on the Mn^2+^ sensitivity phenotype of the Δ*mntP* mutant (Fig. 4C). We noted that the Mn^2+^ sensitivity of the Δ*mntP* mutant (RAS32 strain) was exacerbated by elevating the concentration of MnCl_2_ in the media or increasing the pH of the media. This is expected since increasing concentration of MnCl_2_ in the media correlates with an increase in the cytoplasmic [Mn^2+^] in the Δ*mntP* mutant (Martin et al., 2015; Waters et al., 2011), and alkalinization of the media likewise increases in the cytoplasmic Mn^2+^ (Fig. 4B), leading to Mn^2+^ toxicity. Surprisingly, we noted a pH-dependent boost in the ability of P*_trc_*-expressed Alx to rescue the growth of the Δ*mntP* mutant in media with elevated [Mn^2+^] (Fig. 4C). These results support the notion that the Mn^2+^ export activity of Alx is stimulated by alkaline pH. Alx appears to be a low-activity Mn^2+^ exporter because its rescue ability dropped off at particularly high concentrations of Mn^2+^ (see 350 and 500 µM MnCl_2_ panels), in contrast to MntP. The growth of the Δ*alx* Δ*mntP* double mutant closely resembled that of the Δ*mntP* mutant at elevated media [Mn^2+^] and pH (Fig. 4C). Likewise, the rescue of the growth of the Δ*alx* Δ*mntP* double mutant by overexpression of Alx was similar to that in the Δ*mntP* mutant (Fig. 4C). These observations suggest that chromosomally encoded Alx mitigates the mild perturbations of Mn^2+^ levels brought about by alkaline pH, and its Mn^2+^ export activity appears to be milder in comparison to the chromosomally encoded MntP.

### *alx* expression is post-transcriptionally autoregulated by Mn^2+^ in alkaline pH

As described above, the *alx* translational reporter activity significantly increased at alkaline pH in the Δ*alx* strain (RAS31, Fig. 2B). Intriguingly, this induction was reverted by 1) expressing Alx from a heterologous promoter in the Δ*alx* strain (Fig. 5A) or 2) preserving the chromosomally encoded Alx (Fig. 5B). These results suggest that Alx represses its own expression post-transcriptionally in alkaline pH. Similarly, a 2-fold reduction in the induction of *alx* translational reporter fusion was observed in a strain expressing Alx chromosomally from the native promoter in LB (pH 6.8) supplemented with 500 µM MnCl_2_ (Fig. S5).

**Figure 5.**
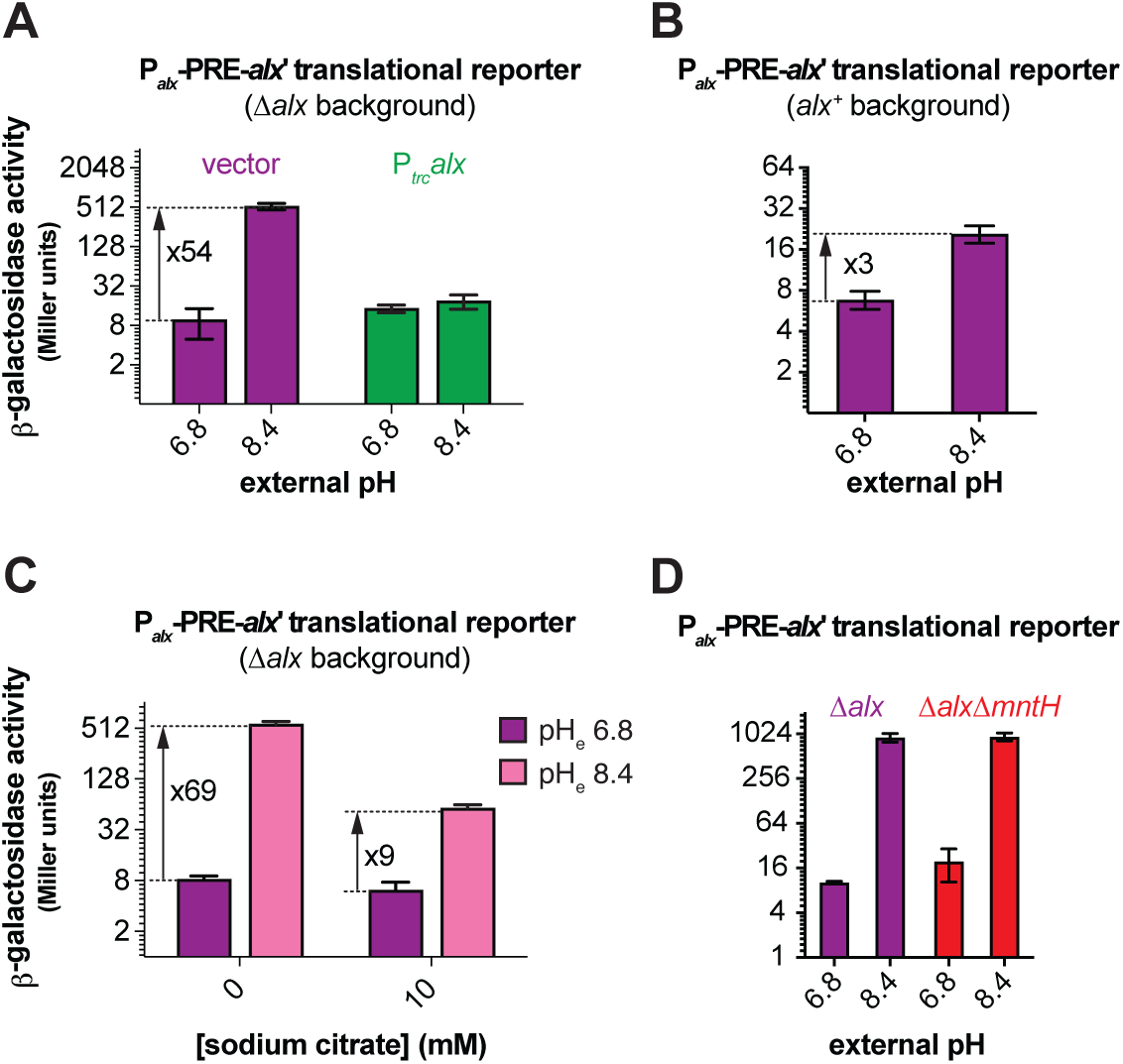
The induction of *alx* expression in alkaline pH and its dependence on [Mn^2+^]. (A) β-galactosidase activity (Miller units) as a reporter of *alx* translation (P*_alx_*-PRE-*alx’-lacZ,* pRA54) was measured in mid-log phase grown cultures of *Δalx*:: Kan derivative (RAS31) bearing an empty vector (pHYD5001) or pHYD5001 expressing Alx from `a` P*_trc_* promoter (pRA27). The cultures were cultivated in LBK media with pHs 6.8 and 8.4, supplemented with appropriate concentration of ampicillin, MnCl_2_ `and` IPTG. (B) β-galactosidase activity (Miller units) as `a` reporter of *alx* translation (P*_alx_*-PRE-*alx’-lacZ*, PRA54) was measured in mid-log phase grown cultures of *alx+* strain (MC4100) in LBK media with pHs 6.8 and 8.4. (C) β-galactosidase activity (Miller units) as a reporter of *alx* translation (P*_alx_*-PRE-*alx’-lacZ*, PRA54) was measured in mid-log phase grown cultures of *Δalx*::Kan derivative (RAS31) in LBK media with pHs 6.8 and 8.4 supplemented with 10 mM sodium citrate. (D) β-galactosidase activity (Miller units) as a reporter of *alx* translation (P*_alx_*-PRE-*alx’-lacZ,* pRA54) was measured in mid-log phase grown cultures of *Δalx* mutant (RAS40) and its *ΔmntH*::Kan derivative (RAS93) in LBK media with pH 6.8 and 8.4. The error shown is standard deviation of three repeats of the experiment.

We noted a 1.5-fold increase in the total Mn^2+^ content in the Δ*alx* strain cultivated in LBK media at pH 8.4 compared to pH 6.8, without any additional Mn^2+^ in the media (Fig. 4B), suggesting that trace Mn^2+^ in LBK was imported into the cells upon alkalinization. These observations offer a new hypothesis where expression of *alx* is induced by elevated intracellular Mn^2+^ brought about by alkaline pH, rather than by alkaline pH directly. To probe this hypothesis, we measured *alx* expression in the Δ*alx* strain in LBK media at pH 6.8 or 8.4 supplemented with 10 mM sodium citrate (Fig. 5C). Citrate chelates divalent and trivalent metal ions in the media, precluding their import into the cells (Anjem et al., 2009; Tong and Rouault, 2007; Westergaard et al., 2017). We noted a significant drop from 69- to 9-fold in the induction of *alx* translation by alkaline pH. These results indicate that elevating intracellular [Mn^2+^] is one of the routes through which alkaline pH induces *alx* expression. The residual 9-fold increase in *alx* translation could be due to incomplete Mn^2+^ chelation by citrate (although 10 mM citrate should be sufficient to chelate 0.25 µM Mn^2+^ in commercial LB broth (Anjem et al., 2009)), an actual direct effect of alkaline pH, and/or effect of other metal ions on *alx* expression. Citrate chelates many divalent metal ions besides Mn^2+^, so this experiment did not completely rule out the involvement of other divalent metal ions in perturbing *alx* expression at alkaline pH. The potential contribution of divalent metal ions other than Mn^2+^ to *alx* expression will be the subject of future work.

One possible mechanism by which alkaline pH may increase the intracellular concentration of Mn^2+^, thereby increasing *alx* translational reporter activity, is by enhancing the activity of MntH, a known transporter for Mn^2+^ uptake in *E. coli*. To address this possibility, we measured the *alx* translational reporter activity in a strain that lacks both chromosomally encoded Alx and MntH (RAS93). The absence of MntH had no impact on the pH-induced increase in the activity of *alx* translation reporter, as the fold induction was the same in the Δ*alx* Δ*mntH* double mutant as in the Δ*alx* mutant (Fig. 5D). These observations indicated that the induction of *alx* translational reporter in alkaline pH is independent of Mn^2+^ uptake by MntH. Similarly, we measured the activity of *alx* translation in LB (pH 6.8) media supplemented with 500 µM MnCl_2_ (Fig. S5). We did not observe any change in the fold induction of the activity of *alx* translation reporter in the Δ*alx* Δ*mntH* double mutant in comparison to that of the Δ*alx* mutant. These data implicate the presence of an MntH-independent route for Mn^2+^ uptake.

### A set of acidic residues in the transmembrane helices is crucial for Alx-mediated Mn^2+^ export

Currently, no experimental three-dimensional (3D) structural information for either Alx or MntP exists. To glean some insight into Alx architecture, its two-dimensional (2D) topology prediction was obtained with multiple web-based tools listed in Table S4. This prediction identified nine Alx transmembrane segments (TMS1-9, Fig. 6A and Table S4), with an overall N-out (periplasmic) and C-in (cytoplasmic) Alx topology. The predicted length of each TMS is similar (∼20 amino acids), although there are some differences in the lengths of TMS3, TMS6 and TMS7 depending on the tool used and in the localization of amino acid residues (in the periplasmic or cytoplasmic environment) residing near the ends of each TMS. Regardless of the prediction tool used, we noted the presence of acidic residues in the transmembrane segments, which is unusual and may suggest the functional importance of these side chains, as demonstrated previously for the export of Mn^2+^ by MntP (Zeinert et al., 2018) (Fig. 6A and Table S4). Two of these residues (D92 and D222) were found to be conserved across members of the TerC family to which Alx belongs (Zeinert et al., 2018). To complement 2D prediction, we also examined an AlphaFold-predicted 3D structure of Alx. Similar to topology predicted arrangement, Alx displayed an N-out C-in conformation in the structure predicted by the AlphaFold server (Fig. 6B). The major difference was in the length of TMS3 in AlphaFold (69 to 103 amino acids) vs topology predicted structure (88 to 101 amino acids (Table S4)). Additional acidic residues localized to the TMS regions of Alx structure predicted by the AlphaFold, e.g., D73 and E86 in TMS3 and D222 in TMS6. The Alx structure predicted by the AlphaFold server was in complete agreement with the RoseTTAFold server prediction (data not shown).

**Figure 6.**
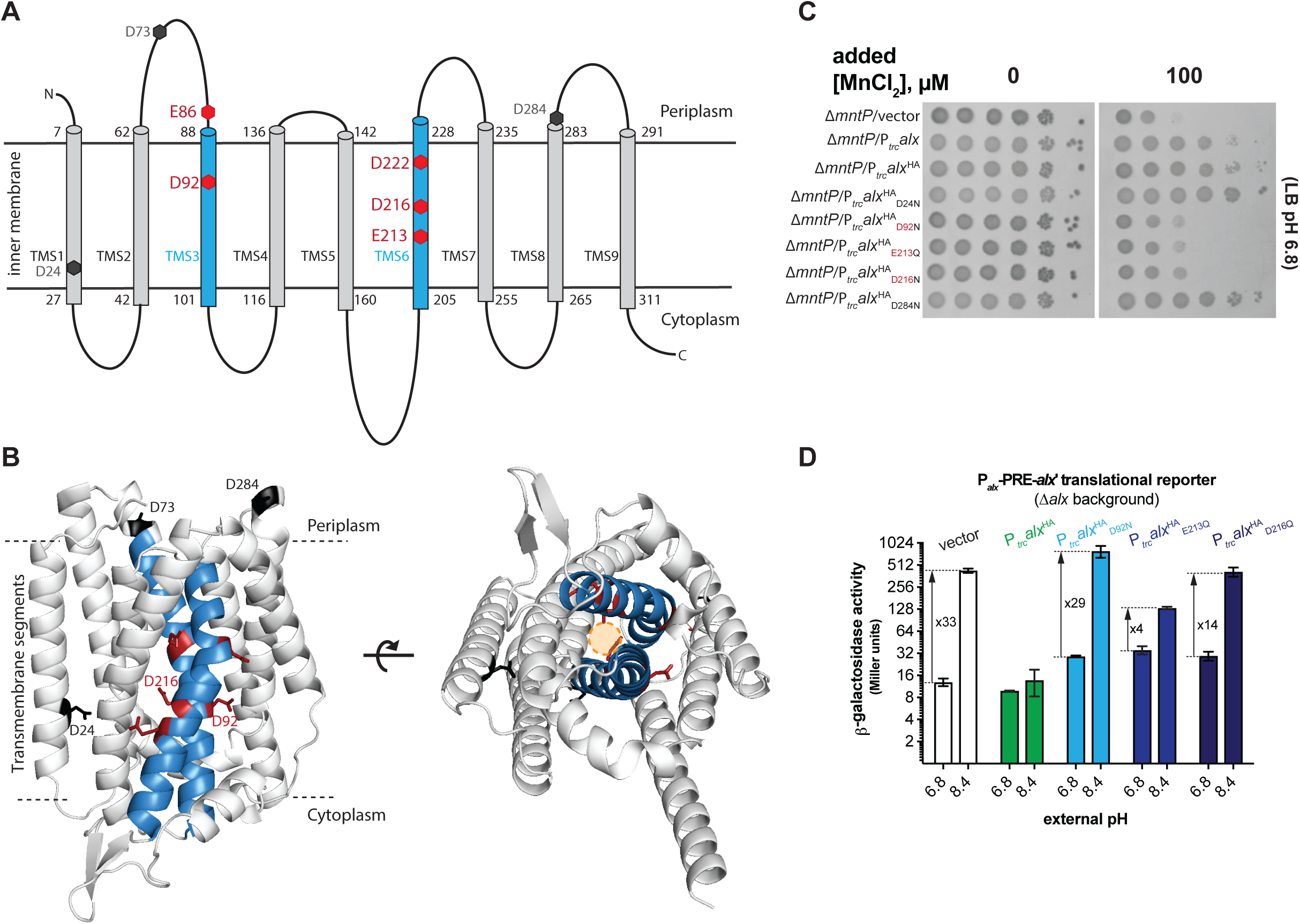
Structural model of Alx and functional relevance of its negatively charged residues in Mn^2+^ export. (A) The two-dimensional topological model of Alx structure predicted with DeepTMHMM algorithm and relative positions of negatively charged residues in transmembrane segments (TMS). (B) AlphaFold-predicted structure of Alx and relative positions negatively charged residues in TMS. A hypothetical path for the export of Mn^2+^ is displayed in the predicted structure. (C) Tenfold serial dilutions of overnight-grown cultures of *ΔmntP*::Kan mutant bearing one of the following plasmids were spotted on the surface of LB agar containing the appropriate concentration of ampicillin, MnCl_2_ and IPTG: a vector (pHYD5001), a derivative of pHYD5001 expressing Alx from a P*_trc_* promoter (pRA27), a derivative of pHYD5001 expressing Alx_HA_ from a P*_trc_* promoter (pRA50), a derivative of PRA50 expressing Alx^HA^_D24N_ (PRA61), Alx^HA^_D92N_ (PRA62), Alx^HA^_E213Q_ (PRA63), Alx^HA^_D216N_ (PRA64), and Alx^HA^_D284N_ (PRA58). (D) β-galactosidase activity (Miller units) as a reporter of *alx* translation (P*_alx_*-PRE-*alx’-lacZ*, pRA54) was measured in mid-log phase grown cultures of Δ*alx*::Kan strain (RAS31) bearing vector (pHYD5001) and a derivative of pHYD5001 expressing Alx^HA^ (PRA27), Alx^HA^_D92N_ (PRA62), Alx^HA^_E213Q_ (PRA63), Alx^HA^_D216N_ (PRA64) from a P*_trc_* promoter. The cultures were grown in LBK media with pH 6.8 and 8.4, supplemented with appropriate concentration of ampicillin and IPTG. The error shown is standard deviation of three repeats of the experiment.

To test the role of the acidic residues predicted to be in the TMS of Alx, a strategy was employed where the effect of P*_trc_*-driven expression of an HA-tagged Alx bearing conservative (D to N or E to Q) replacements was tested on the growth of the Δ*mntP* mutant in LB media (pH 7.2) supplemented with MnCl_2_. With this strategy, expression of the wild-type HA-tagged Alx rescued the growth of the Δ*mntP* mutant like the tag-less version of Alx, suggesting that the HA tag did not alter the activity of Alx. The P*_trc_*-driven expression of Alx bearing E86Q, D92N, E213Q, D216N, or D222N replacement (denoted as Alx^HA^_E86Q_, Alx^HA^_D92N_, Alx^HA^_E213Q_, Alx^HA^_D216N_, or, Alx^HA^_D222N_ respectively) did not rescue the growth of the Δ*mntP* mutant in media supplemented with MnCl_2_, whereas Alx bearing D24N, D73N or D284N substitution did so (Figs. 6C and S6). The expression of HA-tagged Alx mutants was unchanged in comparison to the wild-type Alx (Fig. S7), ruling out Alx expression defects as a cause for the failure to rescue the Δ*mntP* mutant. These results indicated that acidic residues E86, D92, E213, D216 and D222 are important for Alx-mediated Mn^2+^ export.

To experimentally determine compartment-specific positions of Alx acidic residues mentioned above, we followed the substituted cysteine accessibility method (SCAM) as described previously (Butler et al., 2013; Dubey et al., 2021). We substituted residues of interest with a cysteine (Cys, one at a time) and then probed the accessibility of this Cys toward methoxypolyethylene glycol maleimide (Mal-PEG). A DNA sequence encoding N-terminally HA-tagged Alx with an acidic residue-to-Cys mutation was cloned into a plasmid pHYD5001. A culture of the Δ*alx* mutant expressing Cys Alx^HA^ mutant was first separately treated with *N*-ethylmaleimide (NEM) or sodium (2-sulfonatoethyl)methanethiosulfonate (MTSES), to block solvent-exposed Cys residues and prevent their further reaction with Mal-PEG. NEM is membrane permeable and blocks both cytoplasmic and periplasmic Cys on proteins, whereas MTSES is impermeable to the inner membrane and therefore blocks only periplasmic Cys. A Cys that is not blocked by these reagents forms a covalent adduct with the maleimide moiety of Mal-PEG, producing an ∼5 kDa shift in protein mobility on an SDS-PAGE gel. Thus, the reactivity of a particular Cys substitution toward Mal-PEG can infer the topological location of that substituted residue. After blocking the samples with NEM and MTSES, cells were washed, lysed, and then labelled with Mal-PEG. First, we validated the method by investigating the D284 position, which is expected to localize to the periplasmic face of the Alx protein at the end of TMS7 (Fig. 6A). As expected, the D284C mutation did not display reactivity toward Mal-PEG in the presence of MTSES (Fig. S9). Next, we investigated the compartment-specific positions of residues 92, 213, and 216, all of which are predicted to be buried in the inner membrane of the Alx protein (Figs. 6A and B). We observed that Mal-PEG shifted the mobility of Alx in the case of E213C substitution treated with NEM and MTSES (Fig. S9), indicating that E213 is buried in the membrane, as predicted by topology and AlphaFold models. In the case of D92C, Mal-PEG shifted the molecular weight of Alx in the presence of MTSES; however, the bulk of Alx did not display a shift in mobility in the presence of NEM (Fig. S9) suggesting that either D92 is localized to an inner membrane region with solvent accessibility or D92 is in very close proximity to the cytoplasm. Similarly, in the case of D216C, NEM partially blocked the cysteine residues. Thus, Mal-PEG did not shift the molecular weight of Alx completely in the presence of NEM (Fig. S9), suggesting that this residue has limited exposure to the cytoplasmic environment.

To expand upon the functional role of inner membrane acidic residues in the Alx protein, we performed a suite of *alx* translational reporter fusions. Our prior reporter assays demonstrated that Alx displays negative autoregulation, since a translational reporter of *alx* was not induced by alkaline pH or elevated [Mn^2+^] if Alx was expressed from P*_trc_*, presumably because of Alx-mediated Mn^2+^ export (Figs. 5A, 2D and S5). We thus took advantage of this behavior and employed Alx mutants defective in Mn^2+^ export to link the effects of alkaline pH and elevated Mn^2+^ on *alx* expression. Specifically, we tested the impact of P*_trc_*-driven expression of Alx^HA^_D92N_, Alx^HA^_E213Q_ and Alx^HA^_D216N_ on *alx* translational reporter activity in LBK pH 6.8 or 8.4 (Fig. 6D) and LB (pH 6.8) supplemented with 500 µM MnCl_2_ (Fig. S8). Expression of wild-type Alx^HA^ from a plasmid repressed the activity of *alx* translational reporter at alkaline pH in the Δ*alx* strain (RAS31) as expected. The reason behind the lower (33-vs 68-fold) pH-induced increase in *alx* translation in this experiment (strain co-transformed with the translational reporter and Alx expression vector) vs earlier experiment (Fig. 2B, strain transformed with translational reporter only) is unclear. Nevertheless, expression of either Alx^HA^_D92N_, Alx^HA^_E213Q_ or Alx^HA^_D216N_ did *not* repress the activity of *alx* translational reporter activity as wild-type Alx^HA^ did. The fold induction of *alx* translational reporter activity varied (29, 4 and 14 for Alx^HA^_D92N_, Alx^HA^_E213Q_ and Alx^HA^_D216N_, respectively), indicating that Alx_HAE213Q_ and Alx^HA^_D216N_ retain partial activity, whereas Alx^HA^_D92N_ is inactive. We likewise observed the inability of Alx_HAD92N_, Alx^HA^_E213Q_ and Alx^HA^_D216N_ to repress Mn_2+_-induced increase in *alx* translational reporter activity (Fig. S6). However, because this fold increase (∼2) is significantly lower than in alkaline pH, no conclusions about the extent of mutant Alx activity loss could be drawn. Overall, negative autoregulation of *alx* expression in response to alkaline pH is no longer observed when Alx mutants defective in Mn^2+^ transport are expressed; in other words, *alx* expression stays “on”. This suggests a connection between the induction of *alx* expression and Alx-mediated export of Mn^2+^ in alkaline pH, where the return of intracellular Mn^2+^ back to its “healthy” levels via Alx export shuts down further production of Alx at alkaline pH.

### Alx is monomeric *in vivo*

To examine the oligomeric state of Alx and MntP, we employed a non-cleavable membrane permeable crosslinker, disuccinimidyl suberate (DSS). Two N-hydroxysuccinimide (NHS) esters of DSS react with primary amines on proteins (lysines and the N-terminus) resulting in a protein mobility shift on an SDS-PAGE gel. The crude membrane fractions of the Δ*alx* or Δ*mntP* strain expressing N-terminally HA-tagged Alx or MntP, respectively, from a plasmid were isolated and treated with DSS or DMSO as a solvent control (described in Materials and Methods). The samples were electrophoresed by SDS-PAGE and immunoblotted against the HA tag. We did not observe a mobility shift for either Alx or MntP (Fig. 7A), whereas a positive control of an HA-tagged MscL, a mechanosensitive channel, expressed in a Δ*mscL* strain displayed multiple oligomeric forms, consistent with earlier reports (Blount et al., 1996; Dubey et al., 2021; Pathania et al., 2016). These results indicated that Alx and MntP likely exist in a monomeric form *in vivo*.

**Figure 7.**
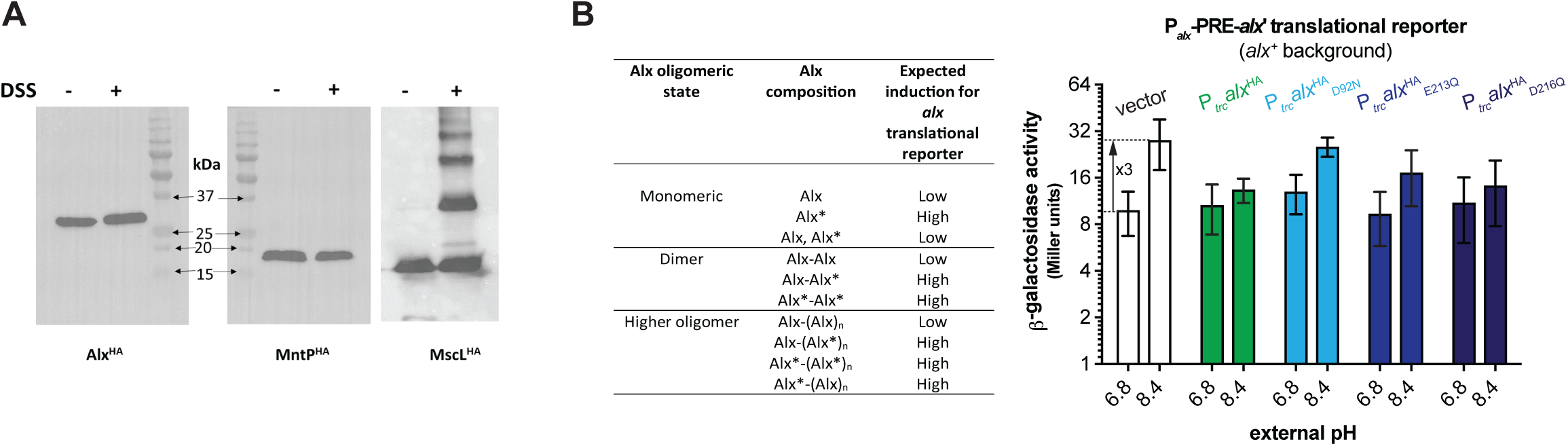
Evidence for a monomeric state of Alx and MntP *in vivo.* (A) Immunoblots probed with anti-HA antibody after electrophoretic mobility of samples containing crude membrane preparations. The crude membranes were harvested for strains RAS31, RAS32, and RAS130 expressing Alx^HA^, MntP^HA^, and MscL^HA^, respectively, and treated with a crosslinker DSS. (B) β-galactosidase activity (Miller units) as a reporter of *alx* translation (P*_alx_*-PRE*-alx’-lacZ,* pRA54) was measured in mid-log phase grown cultures of *alx*^+^ strain (MC4100) bearing a vector (pHYD5001) or a derivative of pHYD5001 expressing Alx^HA^ (PRA27), Alx^HA^_D92N_ (PRA62), Alx^HA^_E213Q_ (PRA63), or Alx^HA^_D216N_ (PRA64) from a P*_trc_* promoter. The cultures were grown in LBK media with pH 6.8 and 8.4, supplemented with appropriate concentration of ampicillin and IPTG. The expected outcome of the experiment is summarized in the table. The error shown is standard deviation of three repeats of the experiment.

A limitation of the DSS crosslinking experiment above is that it relies on the spatial proximity of two primary amines to form the crosslink between protein monomers. An alternative experiment was thus performed to further support Alx’s existence as a monomer *in vivo*. Specifically, we used Alx mutants defective in Mn^2+^ transport as a tool. As described in an earlier section, expression of Alx^HA^_D92N_, Alx^HA^_E213Q_ or Alx^HA^_D216N_ from a synthetic promoter 1) did not rescue the growth of the Δ*mntP* mutant in LB media supplemented with 100 µM MnCl_2_ (Fig. 6C) and 2) increased the signal from *alx* translational reporter, in contrast to the decrease in the reporter output upon expression of the wild-type Alx from either a plasmid or the chromosome (Fig. 6D). If Alx is dimeric or multimeric *in vivo*, then co-expression of the active wild-type Alx from the chromosome and inactive Alx mutants from a plasmid would induce the activity of *alx* translational reporter. Otherwise, *alx* translational reporter activity will remain repressed if Alx is monomeric *in vivo*. In other words, the P*_trc_*-driven expression of an Alx mutant that is defective in Mn^2+^ transport will “poison” the activity of Alx expressed from its native promoter on the chromosome if complexes of these proteins are dimeric or form higher-order oligomers. The effect of P*_trc_*-driven expression of Alx^HA^_D92N_, Alx^HA^_E213Q_ or Alx^HA^_D216N_ was tested on *alx* translational reporter fusion in a wild-type strain (MC4100) that encodes Alx chromosomally. The P*_trc_*-driven expression of Alx and its mutated derivatives left the activity of *alx* translational reporter repressed in pH 8.4 media (Fig. 7B). These data are indicative of the monomeric Alx state *in vivo*, consistent with the DSS crosslinking results.

## Discussion

In this work, we investigated in depth the effect of extracellular alkaline pH and elevated concentration of Mn^2+^ on *alx* expression and provided evidence for Alx export of Mn^2+^ upon alkalinization of the cytoplasm. Our results with *alx* transcriptional and translational reporters corroborate earlier findings that *alx* expression is regulated by both alkaline pH and elevated [Mn^2+^] (Bingham et al., 1990; Dambach et al., 2015; Nechooshtan et al., 2009; Stancik et al., 2002) in a riboswitch-dependent manner. Using a pH responsive GFP variant (pHluorin, (Martinez et al., 2012)), we confirmed that cytoplasm indeed alkalinizes when cells are grown in media with alkaline pH; therefore, our observed changes in gene expression and intracellular metal ion content are a consequence of alkaline cytoplasmic pH. The absence of Alx had no impact on cytoplasmic alkalinization with increasing media pH (Fig. 3B) and did not affect cellular growth in alkaline media (Fig. 3A), ruling out direct Alx involvement in pH homeostasis. Expression of Alx did, however, lower intracellular [Mn^2+^] (and not other tested transition metal ions), but only at alkaline pH (Fig. 4B), thus implicating Alx as a Mn^2+^ exporter in alkaline pH. With this newly uncovered function of Alx, our work points to a connection between the two environmental cues: alkaline pH and elevated [Mn^2+^]. A recent study demonstrated that cytosol alkalinizes in the presence of excess extracellular Mn^2+^ due to an increase in ammonia production within an *E. coli* cell (Kalita et al., 2022); here, we show that the reverse is also true: an alkaline environment promotes the import of Mn^2+^ into the cell.

### Intracellular pH and Mn^2**+**^ content are linked

We find that alkalinization of the cytoplasm leads to an increase in the intracellular concentration of Mn^2+^. Specifically, our intracellular metal ion measurements show a 1.5-fold increase in [Mn^2+^] at pH 8.4 vs 6.8 in the Δ*alx* strain, from 28 to 42 µM (Fig. 4B). Additional indirect pieces of data support this increase. First, the Mn^2+^ sensitivity of Δ*mntP* mutant is exacerbated at alkaline pH (Fig. 4C). Second, even though alkaline pH alone did not impact *mntP* translation, a combination of alkaline pH and extra Mn^2+^ in the media led to a greater *mntP* induction than Mn^2+^ alone (18 and 11-fold, respectively, Fig. 2E). Because upregulation of *mntP* translation is directly proportional to [Mn^2+^] (Fig. 2D), the additional increase is likely due to the additional Mn^2+^ imported from the medium into the cell at alkaline vs. neutral pH. The fact that alkaline pH alone had no effect on *mntP* translation suggests that there is a threshold intracellular [Mn^2+^] needed to begin producing additional MntP (>42 µM). Third, the introduction of a metal chelator (citrate) into the growth medium significantly reduced the alkaline pH-induced increase in *alx* translation, from 68- to 9-fold (Fig. 5C). Citrate is unlikely to be ingested by *E. coli* to affect the expression of *alx* (Ingolia and Koshland, 1979); therefore, the observed drop in *alx* induction is likely due to citrate chelating trace Mn^2+^ in the medium and preventing it from being imported. The mechanism of the alkaline pH-induced Mn^2+^ import is unclear at this point but does not involve the only characterized Mn^2+^ importer MntH in *E. coli* K12 (Figs. 5D and S5), implicating a potential alternative path for Mn^2+^ into the cell.

Why would a cell import Mn^2+^ upon cytosol alkalinization? Mn^2+^ acts as a redox center in the superoxide dismutase SodA and other mononuclear metal enzymes where it can replace Fe^2+^ as a cofactor to prevent protein damage in response to oxidative stress (Anjem et al., 2009; Hopkin et al., 1992; Whittaker et al., 2006). It may thus be an adaptive strategy that cells import Mn^2+^ in response to elevated ROS in alkaline pH. To test this hypothesis, our study employed the *katG* transcriptional reporter to measure the oxidative stress in the parent strain and its Δ*alx* derivative. The readout was indicative of mild oxidative stress in alkaline pH (Fig. 3C). This mild oxidative stress in the parent strain was marginally repressed by supplementation of Mn^2+^ or by the absence of chromosomally encoded Alx (Fig. 3D). An increase in the activity of Mn^2+^-dependent SodA is likely behind the repression of mild oxidative stress by supplemented Mn^2+^ (Pugh et al., 1984), whereas accumulation of Mn^2+^ in the Δ*alx* strain (due to lack of Alx-mediated Mn^2+^ export) is likely responsible for oxidative stress repression in the Δ*alx* strain. These observations support our finding that Alx exports Mn^2+^ and is thereby indirectly related to redox homeostasis. Nevertheless, the physiological basis for mild induction of oxidative stress in alkaline pH and its relationship to *alx* expression is still unclear.

The heterologous expression of MntP (Mn^2+^ exporter) and Alx slowed the aerobic growth of a sensitized strain that lacks hydrogen peroxide degrading enzymes (AhpCF and KatG) (Zeinert et al., 2018). The effects of MntP overproduction on the growth of Δ*ahpCF* Δ*katG* double mutant were explained by reduced intracellular [Mn^2+^] and corresponding reduced protection from elevated ROS, believed to occur through reduced activity of Mn^2+^-dependent SodA and reduced protection of mononuclear enzymes where Fe^2+^ acts as a cofactor (Anjem et al., 2009). However, similar effects of Alx overproduction on the growth of the abovementioned strain were explained differently by (Zeinert et al., 2018) as Alx was viewed as an importer of Mn^2+^. Alx export of Mn^2+^ by analogy to MntP, on the other hand, better explains the observed slower growth of the Δ*ahpCF* Δ*katG* strain upon Alx overexpression because P*_trc_*-driven expression of Alx rescues the growth of the Δ*mntP* mutant in media supplemented with extra Mn^2+^ and reduces the intracellular Mn^2+^ levels in the alkaline pH.

The *alx* translational reporter displayed a 68-fold induction in alkaline pH media and a 22-fold induction in neutral pH media containing 500 µM MnCl_2_, with an 86-fold induction when two environmental cues (alkalinity and high [Mn^2+^]) were combined (Fig. 2). These results favor the notion that alkaline pH augments the effects of elevated [Mn^2+^] on *alx* expression. The alkaline pH-induced Mn^2+^ import also provides an alternate explanation to a recent report that elevated cytoplasmic [Mn^2+^] results in higher activation of *mntP* riboswitch upon alkalinization in media supplemented with excess Mn^2+^, in contrast to the proposed tighter interaction between Mn^2+^ and riboswitch element (Kalita et al., 2022) Differences in the fold induction by the two cues are reflective of a distinct mechanism that may operate for modulation of *alx* expression in alkaline pH that depends on elevated cytoplasmic [Mn^2+^]. One of the future directions for deconvoluting the mechanism of pH and Mn^2+^ control of *alx* expression will be to examine how pH and Mn^2+^ differentially affect *alx* mRNA folding, and specifically folding of its 5’ UTR riboswitch. The second direction will focus on the identification of novel pH-dependent transporters that address the involvement of other metal ions besides Mn^2+^ at alkaline pH to enhance *alx* expression.

### Mn^2+^ export by Alx

A previous study proposed that Alx may function as an Mn^2+^ uptake protein based on the cellular [Mn^2+^] measurements in the presence of supplemented Mn^2+^ in the media (Zeinert et al., 2018). Contrary to earlier studies, here we provided multiple pieces of evidence for the Alx-mediated *export* of Mn^2+^ in alkaline pH or conditions where cytoplasmic Mn^2+^ levels are elevated. First, the inability of Δ*mntP* mutant to grow in the media with elevated [Mn^2+^] was partially rescued with P*_trc_*-driven expression of Alx (Fig. 4A). This rescue phenotype was missed in the previous work (Zeinert et al., 2018) likely because of the following differences between our and this prior experimental setup: (*i*) Alx was expressed from a weaker promoter (P_BAD_ vs stronger P*_trc_* in our work) and (*ii*) rescue experiments were performed at neutral pH only. Strikingly, the rescue of Δ*mntP* mutant’s sensitivity towards Mn^2+^ by P*_trc_*-driven expression of Alx becomes more and more pronounced with increasing pH, while Mn^2+^ sensitivity of Δ*mntP* mutant becomes exacerbated with increasing pH (Fig. 4C). Secondly, the P*_trc_*-driven expression of Alx in the Δ*alx* mutant resulted in ∼2-fold reduction of intracellular [Mn^2+^] but only in alkaline pH, returning intracellular [Mn^2+^] from 42 to 19 µM (Fig. 4B). Therefore, alleviation of the growth of the Δ*mntP* mutant in media with alkaline pH and extra Mn^2+^ by P*_trc_*-driven expression of Alx can be explained by increased activity of Alx in alkaline pH. Overall, our observations corroborate that the Alx export of Mn^2+^ is stimulated by alkaline pH.

Mn^2+^ export activity of Alx at alkaline pH is also supported by the observed negative feedback regulation of *alx* expression. Specifically, the alkaline induction of *alx* translational reporter (68-fold) was repressed by the presence of Alx encoded chromosomally from a native promoter or expressed from P*_trc_* (Figs. 2B and 5AB). These results can be explained if Alx exports Mn^2+^ thereby reducing cytoplasmic [Mn^2+^] to the levels that no longer stimulate *alx* translation. In another observation, the expression of Alx chromosomally from a native promoter resulted in only a two-fold reduction in the activity of the *alx* translational reporter in the pH 6.8 media supplemented with 500 µM MnCl_2_ (Fig. S5). The mild reporter activity reduction, in this case, can be explained by the lack of alkaline pH-stimulated Mn^2+^ export activity of Alx.

The driving force and mechanism behind Alx’s export of Mn^2+^ remain a mystery to us. A proton gradient is unlikely to drive this transport because we do not observe a loss of pH gradient upon Mn^2+^ introduction to inverted membrane vesicles containing Alx (Fig. S4). This observation rules out the possibility that Alx is an Mn^2+^/H^+^ antiporter. In another system, a high concentration of potassium ion (K^+^) in the media was proposed to stimulate the activity of K^+^ export proteins and inhibit the activity of K^+^ uptake proteins (Li et al., 2002; Roe et al., 2000; Sharma et al., 2016). However, in the case of Alx, the stimulation of its activity by alkaline pH is unlikely through just an increase in cellular Mn^2+^ levels. This reasoning was supported by the observations that the rescue of Mn^2+^ sensitivity of Δ*mntP* mutant by P*_trc_*-driven expression of Alx is absent in the media with high [Mn^2+^] in comparison to intermediate [Mn^2+^] where P*_trc_*-driven expression of Alx rescues the growth of the Δ*mntP* mutant (Fig. 4A, 4C). The most likely explanation for the stimulation of Alx activity in alkaline pH could be pH-driven structural changes in the Alx protein that affect its Mn^2+^ export. We identified acidic residues of Alx (E86, D92, E213, D126, and D222) in TMS3 and TMS6 that are crucial for Mn^2+^ transport, by analogy to MntP and MntH (Fig. 6B, (Haemig and Brooker, 2004; Zeinert et al., 2018)). The interaction of positively charged solute (Mn^2+^) and acidic side chains of TMS3 and TMS6 of Alx may provide a path for Mn^2+^ transport as depicted in Fig. 6B. Some of the key side chains point away from each other in the Alphafold structure of Alx. Alkaline pH-induced conformational changes, however, may reorient TMS 3 and 6 of Alx to allow Mn^2+^ coordination. Such pH-induced conformational change was hypothesized to be the structural basis of alkaline pH-stimulated transport by an *E. coli* Na^+^/H^+^ antiporter, NhaA (Hunte et al., 2005; Taglicht et al., 1991). We speculate alkaline pH-induced conformational changes in the crossed TMS3 and TMS6 of Alx, similar to NhaA (Hunte et al., 2005), where acidic or neutral cytoplasmic pH favors a locked conformation limiting Mn^2+^ export into the periplasm and alkaline pH orients the two helices into an open conformation stimulating the export of Mn^2+^.

A Mn^2+^/Ca^2+^ exporter MgtA in *Streptococcus pneumonia* and a Mn^2+^ exporter YoaB in *Lactobacillus lactis* are putative P-type ATPases that use ATP hydrolysis to translocate their metal cargo against its concentration gradient. The expression of these exporters is regulated by the *yybP*-*ykoY* family of riboswitches (Martin et al., 2019; Price et al., 2015). It would be fascinating to find out whether Alx and MntP, whose expression is likewise regulated by riboswitches from the *yybP*-*ykoY* family, employ active transport to mitigate toxic levels of Mn^2+^ like classical P-type ATPases. However, the cytoplasmic domain of Alx in the AlphaFold structure is much smaller in comparison to P-type ATPases (Farley, 2011). Interestingly, the cytoplasmic loop of Alx contains several positively charged residues that may participate in the binding of ATP, perhaps in concert with a protein partner in the cell and help in the transfer of high-energy acyl phosphate to an aspartate to drive a conformational change needed for Mn^2+^ translocation across the membrane. In our experiments, we did not observe any growth phenotype in the Δ*alx* mutant indicating that chromosomally encoded Alx does not provide a significant advantage in the tested laboratory conditions of additional Mn^2+^ in the media and/or alkaline pH. This observation also suggests that a pH-driven increase in intracellular [Mn^2+^] to 42 µM is not toxic to *E. coli*. Overexpression of Alx, however, did rescue growth of the Δ*mntP* mutant in media with elevated [Mn^2+^] and alkaline pH, suggesting Alx functions as a weak Mn^2+^ exporter in an alkaline environment, meaning that its rate of Mn^2+^ export is too low to be consequential to *E. coli* growth under tested conditions. Curiously, an earlier study reported that *alx* expression is induced in both neutral and alkaline pH under anaerobic growth conditions (Hayes et al). In anaerobic conditions, the absence of superoxide dismutase enzymes (*sodA* and *sodB* double mutant) does not cause a growth defect (Carlioz et al., 1986), implying that ROS stress is minimal. Thus, a cell no longer needs additional Mn^2+^ during growth in an anaerobic environment and preventing Mn^2+^ buildup due to its uptake becomes important. This explains why *alx* is expressed even at neutral pH in anaerobic conditions (Hayes et al., 2006). We speculate that the expression of *alx* may provide an advantage in environmental niches where *E. coli* and other enterobacteria are challenged by both alkaline pH and hypoxic conditions, such as the human gut (Litvak et al., 2018; Rogers et al., 2021). The elevation of cellular Mn^2+^ levels in alkaline conditions (Fig. 4B) favors the expression of *alx* to get rid of excess Mn^2+^. To cope with the threat of high [Mn^2+^] in the environment, the Mn^2+^ export activity of chromosomally encoded MntP is sufficient to protect the cell. On the other hand, when changes in intracellular [Mn^2+^] are mild, e.g., as brought about by alkaline pH, Alx fulfills the job of maintaining healthy levels of Mn^2+^ inside the cell. We thus pose that Alx-mediated Mn^2+^ export provides a primary protective layer that fine-tunes the cytoplasmic Mn^2+^ levels, especially during alkaline stress.

## Acknowledgements

We would like to thank Drs. Abhijit Sardesai, James Imlay, and Robert Browne for graciously providing plasmids and strains. We thank the members of the Mishanina lab for critical reading and helpful suggestions for improving the manuscript, as well as Drs. Manuel Raffatellu and Daniel Roston. We are grateful to Dr. Neal Arakawa at the Environmental and Complex Analysis Laboratory (ECAL) at UCSD for assistance with ICP-MS measurements and to Dr. Terence Hwa for the pHluorin plasmid and use of the plate reader for intracellular pH and growth rate measurements. We would like to thank Iman Saeed for her assistance with cloning experiments. We also thank Drs. Mark Herzik and Galia Debelouchina for providing access to their lab instruments. This work was funded by the National Institutes of Health/National Institute of General Medical Sciences (ESI; grant no. R35 GM142785), UCSD institutional support, and Yinan Wang Memorial Chancellor’s Endowed Junior Faculty Fellowship to T. V. M.

## Materials

Manganese (II) chloride tetrahydrate and potassium benzoate were purchased from Alfa Aesar. Tris(hydroxymethyl)methyl-3-amino propane sulfonic acid (TAPS) was purchased from Acros Organics. Mal-PEG, N-(1,1-dimethyl-2-hydroxyethyl)-3-amino-2-hydroxy-propane sulfonic acid (AMPSO), nigericin sodium salt and valinomycin were purchased from Sigma-Aldrich. ACMA was purchased from Invitrogen. NEM and o-nitrophenyl-β-D-galactopyranoside were purchased from Thermo Scientific. Isopropyl-β-D-thiogalactopyranoside (IPTG) and 3-(N-morpholino) propane sulfonic acid (MOPS) were purchased from Fisher Scientific. MTSES was purchased from Biotium.

## Methods

### Tests of the Mn^2+^ sensitive phenotype and its rescue

Strains were inoculated in LB broth overnight at 37 °C in a shaker. The 5 µl of 10-fold serial dilutions of an overnight culture of an appropriate strain was spotted on LB agar supplemented with MnCl_2_ as described in the Results. Whenever required, LB broth or agar media were supplemented with an appropriate concentration of antibiotics and IPTG. LB agar plates were imaged after incubation at 37 °C for 14 to 16 hours.

### β-galactosidase assays

Overnight cultures of the strains were inoculated in LBK broth with pH 6.8 and 8.4 or in LB broth with or without appropriate concentration of MnCl_2_ at 37 °C to a mid-log phase. The appropriate concentration of antibiotics (trimethoprim and/or ampicillin) and IPTG (1 mM) were supplemented when needed in the experiments. β-galactosidase assays were carried out by following the method of Miller, and β-galactosidase-specific activity was reported in Miller units (Miller JH, 1992). Each reported value with a standard deviation is the average of three independent experiments.

### ICP-MS measurement of cellular metal ions

The total amounts of Mn^2+^, Fe, and Zn^2+^ were quantified from 5-ml cultures. Cells were grown overnight in LB broth and then inoculated in LBK pH 6.8, and LBK pH 8.4 media supplemented with 1 mM IPTG and appropriate concentration of ampicillin respectively. After growth to the mid-log phase at 37 °C, cells were harvested using centrifugation at 4000*g* for 10 minutes. Cell pellets were washed with 10 mM N-2-hydroxyethylpiperazine-N-2-ethane sulfonic acid (HEPES) pH 7.5, containing 2 mM EDTA and then washed twice with 10 mM HEPES as described in (Zeinert et al., 2018). Cell pellets were dried for 1 hour in a centrifuge evaporator. Dried cell pellets were solubilized in 400 µl of 30% (v/v) HNO_3_ and incubated at 95 °C for 10 min. Then samples were centrifuged at 20,000*g* for 5 minutes. Samples were prepared for inductively coupled plasma mass spectrometry (ICP-MS) by diluting 300 µl of supernatant of lysed cells into 2.7 ml of 2.5% (v/v) HNO_3_ and run on an iCAP RQ ICP-MS (Thermo Scientific). Metal concentrations are presented as intracellular levels after correction for mean cell volume determined from total protein content (Martin et al., 2015). The data obtained were presented from three repeats of the experiment.

### Cytoplasmic pH measurements

The wild-type strains of *Escherichia coli* and its Δ*alx* mutant containing expressing pHluorin (pRA46) were grown overnight in LBK medium buffered with 50 mM of MOPS with pH 7.5 and an appropriate concentration of ampicillin. Cells were inoculated and grown to mid-log phase in fresh LBK medium with pH 7.5 with an appropriate concentration of ampicillin and 1mM IPTG at 37 °C. Cells were harvested from appropriate volume of the cultures by spinning at 4000 rpm. Then cells were resuspended in 4 ml of M63A minimal medium [0.4 g/liter KH_2_PO_4_, 0.4 g/liter KH_2_PO_4_, 2 g/liter (NH_4_)_2_SO_4_, 7.45 g/liter KCl supplemented with 2 g/liter casein hydrolysate) and buffered to the desired pH with 50 mM concentration of the appropriate buffer pHs 7.0 and 7.5, MOPS; 8.5, TAPS; and pHs 9 and 9.5, AMPSO. Due to poor growth in extremely alkaline conditions, the initial *A*_600_ for cells growing in M63A media with external pH (pH_e_) 9 and 9.5 was chosen to be around 0.2, and in M63A medium with pH_e_ 7, 7.5, and 8.5 was chosen to be around 0.05. The cultures were grown for 2 hours at 37 °C with mild shaking. To generate a standard curve, 95 µl volume of the culture of parent strain expressing pHluorin from each buffered media was withdrawn and mixed with potassium benzoate to a final concentration of 40 mM in 96-well plates. The cultures were incubated at room temperature for 3 min. Methanol amine was added to the culture at a final concentration of 20 mM. The cultures were incubated for 3 min at room temperature. The 100 µl of the parent strain and its Δ*alx* mutant expressing pHluorin were withdrawn from each buffered media to 96 well plates and used for the internal pH (pH_i_) measurements. The measurements with fluorescence intensity at 530 nm were taken for the two excitation (410 and 470 nm) wavelengths for each strain expressing pHluorin as described in (Martinez et al., 2012). The ratio of fluorescence intensity of pHluorin at two excitation wavelengths against pH was plotted to generate a standard curve. The slope of the curve was used to calculate the pH_i_ across different pH_e_.

### *In vivo* cross-linking with disuccinimidyl suberate

To determine the oligomeric state of Alx and MntP, we performed *in vivo* cross-linking with disuccinimidyl suberate (DSS). The strains RAS31 (MC4100 Δ*alx*::Kan), RAS32 (MC4100 Δ*mntP*::Kan), and RAS130 (MC4100 Δ*mscL*::Kan) bearing the plasmid pRA50, encoding Alx^HA^, pRA70 encoding MntP^HA^ and the plasmid pRA72 encoding MscL^HA^ (MscL bearing a C-terminal HA tag) respectively, were grown in 100 ml of LB with appropriate concentration of ampicillin. Cells were simultaneously induced with 1 mM IPTG for P*_trc_*-driven expression of Alx^HA^, MntP^HA^, and MscLC^HA^ from the above-mentioned plasmids. Cells were grown to the mid-log phase and normalized for *A*_600_ of 50. Cells were harvested by centrifugation and pellets were washed in reaction buffer (30 mM sodium phosphate, pH 7.5 and 100 mM NaCl), and resuspended in 5 ml of the same buffer as described in (Dubey et al., 2021; Pathania et al., 2016). Cells were broken using a QSONICA sonicator, at 6/10 power 15 seconds on, 15 seconds off for 3 min on ice. The lysate was centrifuged at 12,000 rpm for 15 min to remove cell debris and unbroken cells. The supernatant was subjected to an ultracentrifugation step, at 48,000*g* for 90 min. The recovered crude membrane pellet was resuspended in 4 ml of reaction buffer. Two ml of the membrane suspension was transferred to two tubes. DSS was added into one tube at 1 mM of final concentration and the other received an equal volume of solvent (DMSO). Two tubes were kept on a shaking platform for 30 min at room temperature. The reaction was quenched with the addition of Tris-HCl pH 8 at 100 mM final concentration. The membrane suspension was pelleted by ultracentrifugation at 48,000*g* for 30 min and solubilized in 100 μl of SDS loading buffer. Samples were loaded and separated in 12% SDS-PAGE, and protein detection was performed by immunoblotting with an anti-HA antibody.

