## Supplemental text and figures for "A riboswitch-controlled manganese exporter (Alx) tunes intracellular Mn^2+^ concentration in *E. coli* at alkaline pH"

List of contents:

**Supplemental Materials and Methods**

- Bacterial strains and growth conditions
- Plasmid construction
- Growth rate measurements
- Immunoblotting
- Preparation and storage of inside-out vesicles
- Assay for the detection of substrate-induced proton release in inside-out membrane vesicles
- Substituted cysteine (Cys) accessibility measurements

**Table S1:** List of *E. coli* K12 strains employed in this study.

**Table S2**: Plasmids employed in this study.

**Table S3**: Oligonucleotide primers employed in this study.

**Table S4:** Comparison of predicted transmembrane segments (TMS) of Alx using prediction tools.

**Supplemental References**

#### Supplemental Materials and Methods

##### Bacterial strains and growth conditions

The strains employed in this study are derivatives of *E. coli* K12 and tabulated in Table S1. The MG1655 Δ*mscL*::Kan was constructed by following the procedures of recombineering as described in (Datsenko and Wanner, 2000). The Δ*alx*::Kan, Δ*mntP*::Kan, and Δ*mntH*::Kan mutations were sourced from appropriate strains from the Keio collection (Baba et al., 2006) and introduced into recipient strains by P1 transduction. The gene encoding the Kanamycin resistance determinant was excised upon treatment with plasmid pCP20 in Δ*alx*::Kan (Datsenko and Wanner, 2000). All strains were cultivated in LB agar or broth at 37 °C. The standard pH of LB agar or LB broth employed in this study was 7.2. The pH of LB agar or broth media was mentioned in the experiments where the pH was not adjusted to 7.2. We also employed potassium-modified LB medium (LBK), where equimolar KCl replaced NaCl, buffered with 100 mM N-[Tris(hydroxymethyl)methyl]-3-amino propane sulfonic acid (TAPS) and pH adjusted with KOH to 8.4) (Stancik et al., 2002). In other experiments LB agar was buffered with HEPES, Tris-Cl, and TAPS (50 mM final concentration), and pH was adjusted to 7, 7.8, and 8.2, respectively, before autoclaving. Whenever required appropriate amounts of kanamycin, ampicillin, chloramphenicol, trimethoprim, L-arabinose, and IPTG were supplemented to the media.

##### Plasmid construction

The plasmid pRA40 carrying $P_{alx}$-PRE-*alx'* (transcriptional reporter of *alx*) was constructed by PCR-amplifying MC4100 chromosomal DNA from 473 nucleotides upstream and 53 nucleotides downstream from the translation start site of *alx* (Nechooshtan et al., 2009), using oligos RAV5 and RAV48, and then cloning the PCR product into the PstI and KpnI sites of a single-copy plasmid pMU2385. The plasmid pRA54 (translational reporter of *alx*) was constructed similarly by PCR-amplifying MC4100 chromosomal DNA from 473 nucleotides upstream and 99 nucleotides downstream from the translation start site of *alx* using oligos RAV5 and RAV23 and cloning of PCR product into the PstI and BamHI sites of single-copy plasmid pMU2386, such that the first 33 codons of the ORF are in frame with the 8th codon of *lacZ*. We extended the length of *alx*′ in the translational reporter of *alx* (98 nucleotides downstream from the translation start site), compared to the transcriptional reporter (53 nucleotides downstream from the translation start site) for two reasons: 1) to include the last putative transcriptional pause proximal to the *alx* translation start site (Fig. S1) and 2) to increase the probability of the Alx′-LacZ hybrid protein to localize in the cytoplasm (refer to the topological analysis of Alx in Fig. 6A). The ΔPRE derivatives of transcriptional (pRA41) and translational (pRA55) reporters of *alx* were constructed by site-directed mutagenesis of plasmids pRA40 and pRA54, respectively, using oligos RAV15 and RAV16. Similarly, a derivative of transcriptional (pRA49) and translational (pRA56) reporters of *alx* were constructed where the length of *alx*′ was shortened. The pRA49 was constructed by PCR-amplifying MC4100 chromosomal DNA using oligos RAV5 and RAV117 and cloning the PCR product into the PstI and KpnI sites of pMU2385. Similarly, pRA56 was constructed by PCR-amplifying MC4100 chromosomal DNA using a pair of oligos RAV5 and RAV118 and cloning the PCR product into the PstI and BamHI sites of pMU2386.

The pRA48 plasmid containing transcriptional reporter of *mntP* was constructed by PCR amplifying MC4100 chromosomal DNA from 882 nucleotides upstream and 47 nucleotides downstream from the translation start site of *mntP* (Dambach et al., 2015), using oligos RAV119 and RAV120, and then cloning the PCR product into the PstI and KpnI sites of a single-copy plasmid pMU2385. The plasmid pRA57 (translational reporter of *mntP*) was constructed by PCR-amplifying MC4100 chromosomal DNA from 882 nucleotides upstream and 47 nucleotides downstream from the translation start sites of *mntP* (Dambach et al., 2015) using oligos RAV119 and RAV126 and cloning of the PCR product in the PstI and SalI sites of single-copy plasmid pMU2386, such that the first 47 nucleotides of the *mntP* ORF are in frame with the 8th codon of *lacZ*.

Oligos RAV65 and RAV66 were employed to PCR-amplify the gene encoding pHluorin from a plasmid (pGFPR01) obtained from a strain JLS1105 (Martinez et al., 2012). The PCR product was cloned into pHYD5001 using Gibson assembly, producing pRA46. Oligos RAV11 and RAV12 were used to PCR-amplify *alx* from the MC4100 genomic DNA. The PCR product was then cloned into the NdeI and HindIII sites of pHYD5001. The resulting pHYD5001 expressing *alx* from $P_{trc}$

promoter was denoted as pRA27. It was noted upon sequencing of pRA27 that the NdeI site was mutated but *alx* ORF is still retained. The pRA50 plasmid encoding the HA epitope tagged version of Alx on N-terminus from $P_{trc}$ promoter was constructed by PCR-amplification with oligos RAV139 and RAV12 and cloning the product into the NdeI and HindIII sites of pHYD5001. Similarly, pRA70 and pRA72 were constructed to express the HA epitope tagged MntP and MscL on their N- and C-terminus, respectively, from $P_{trc}$ promoter. Two PCR products were generated by using oligos RAV178 and RAV14 for *mntP* cloning and a set of three oligos RAV179, RAV180, and RAV181 for *mscL* cloning from MC4100 genomic DNA as template. PCR products were cloned into the NdeI and HindIII sites of pHYD5001. Other plasmids pRA58, pRA61, pRA62, pRA63, pRA64, pRA65, pRA66, PRA67, pRA68, pRA69, pRA74, pRA75 and pRA76 were constructed by site directed mutagenesis of plasmid pRA50 by using oligo pairs stated in Table S3.

**Growth rate measurements**

10 µL of log-phase cultures of appropriate strain were diluted to 1 ml of fresh LB and LBK media with pH 6.8 and 8.4. The appropriate concentration of $MnCl_2$ was added to LB and LBK media as described in the experiments. 200 µL of these diluted cultures were grown in wells of honeycomb multi-well plates at 37 °C while shaking. Growth curves were generated using an automated Spark multimode plate reader by Tecan.

**Immunoblotting**

The cultures of appropriate strains were grown to the mid-log phase. Cells were harvested by centrifugation for 20,000*g* for 1 min after OD normalization. Cell pellets of an $A_{600}$ of 1 were solubilized in 200 µL of SDS-PAGE loading dye. The whole cell extracts were loaded on 12% SDS-PAGE gel after incubation at 37 °C for 10 min. Proteins were transferred to the PVDF membrane (Bio-Rad) by Trans-Blot Turbo, a semi-dry transfer apparatus (Bio-Rad). The PVDF membrane was treated with a blocking buffer (Tris-HCl buffer saline with 5% fat-free milk powder) for 30 min. The membrane was probed with an anti-HA rabbit monoclonal antibody (Invitrogen), at a dilution of 1:5,000 overnight at 4 °C and with an anti-rabbit, horseradish peroxidase-conjugated antibody

(Promega) at a dilution of 1:5,000 for 2 hours. The blot was developed using a Clarity Western ECL substrate (Bio-Rad), and the signal was detected by the ChemiDoc imaging system. The PVDF membrane was stained with 0.1% amido black solution to confirm equal loading of samples across the lanes. Following the staining procedure for 15 s, the membrane was destained with a solution of 45% methanol, 45% water, and 10% glacial acetic acid.

**Preparation and storage of inside-out vesicles**

Inside-out vesicles were prepared from the strain RAS42 that lacks *alx* and *mntP*. RAS42 containing an empty vector and its derivatives that express $Alx^{HA}$ or $MntP^{HA}$ from the *trc* promoter were cultivated in 1 L of LB broth with an appropriate concentration of ampicillin at 37 °C. The cultures were grown until they reached an $A_{600}$ of 0.2 and then induced with 1 mM IPTG for $P_{trc}$-driven expression of $Alx^{HA}$ and $MntP^{HA}$ at 18 °C for 12 hours. The inside-out vesicles were prepared using procedures identical to those described in (Verkhovskaya, 2017) with some modifications for the storage of vesicles. Aliquots of the inside-out vesicle preparations were stored in Buffer B (Verkhovskaya, 2017) with 10 % glycerol, frozen in liquid nitrogen, and stored at –80 °C until further use.

**Assay for the detection of substrate-induced proton release in inside-out membrane vesicles**

Kinetic measurements for the experiments were performed as described in the reference (Dubey et al., 2021). Frozen vesicles were thawed on ice, and 40 µL of vesicles from each samples were used for the assays. 40 µL of membrane vesicles were diluted to 2 mL with solution of 50 mM KCl and 10 mM $MgSO_4$. ACMA (10 µM) and valinomycin (0.05 µM) were added at the beginning of the kinetic measurement. The fluorescence measurements of ACMA ($\lambda_{Ex}$ 409 nm, $\lambda_{Em}$ 474 nm) were recorded for 250 s with continuous stirring of the samples. ATP (0.25 mM, final concentration) was added after 50 s to generate the pH gradient across the membrane as estimated by quenching of ACMA's fluorescence. $MnCl_2$ (1 mM, final concentration) was added after 150 s. Any significant change in pH due to substrate-induced proton release was measured by dequenching of ACMA fluorescence. The measurements were terminated by the addition of nigericin (4 µM) after 200 s.

**Substituted cysteine (Cys) accessibility measurements**

The accessibility of Cys residues introduced into $Alx^{HA}$ that does not contain any Cys residues was tested by the substituted cysteine accessibility method (SCAM) (Butler et al., 2013; Dubey et al., 2021). Cultures of the strain RAS31 bearing plasmids encoding Cys substituted derivatives of $Alx^{HA}$ were cultivated in LB broth with 1 mM IPTG to mid-log. Cells were pelleted by centrifugation at 16000*g* for 3 min and washed with Phosphate Buffered Saline (PBS). Harvested cells were suspended in 200 µL of PBS such that cell suspension measured an $A_{600}$ of 0.8. Resuspended cells were divided into four 50 µL aliquots. Two of the aliquots were treated with cysteine sulfhydryl blockers. The remaining aliquots were treated with either N-ethylmaleimide (NEM) sodium (2-sulfonatoethyl) methane thiosulfonate (MTSES), at a final concentration of 5 mM each. The four tubes were placed in the dark on a shaking platform at room temperature. After 1 hour of incubation, cells were washed twice with PBS and resuspended in 50 µL of lysis buffer (15 mM Tris-HCl, pH 7.4, 1% SDS, 6 M urea). The two blocked samples (with NEM or MTSES) and one sample not exposed to either NEM or MTSES were labeled with Mal-PEG (methoxy polyethylene glycol maleimide, molecular weight 5000 dissolved in DMSO) present at 5 mM. One of the four samples was treated with the solvent DMSO (control). Samples were kept in the dark on a shaking platform for 1 hour. Then samples were processed as per reference (Butler et al., 2013) by adding an equal volume of 2X AB buffer. Samples were sonicated briefly before SDS PAGE loading. Following SDS-PAGE, the gels were processed as described in the procedures for immunoblotting.

**Table S1.** List of *E. coli* K12 strains employed in this study[#].

| Strain | Genotype |
|---|---|
| MC4100 | *Δ(argF-lac)U169 rpsL150 relA1spoT1araD139 flbB5301 deoC1 ptsF25* |
| RAS31 | MC4100 Δ*alx*::Kan |
| RAS32 | MC4100 Δ*mntP*::Kan |
| RAS40 | MC4100 Δ*alx* |
| RAS42 | MC4100 Δ*alx* Δ*mntP*::Kan |
| RAS93 | MC4100 Δ*alx* Δ*mntH*::Kan |
| AL441 | *MG1655Δ(lacZ1::cat)1* attλ::[pSJ501*::katG′-lacZ*$^+$ |
| RAS136 | AL441 Δ*alx*::Kan |
| RAS130 | MC4100 Δ*mscL*::Kan |

[#] All strains employed in this study are *E. coli* K12 derivatives. The Δ*alx*::Kan, Δ*mntP*::Kan, and Δ*mntH*::Kan deletion mutations were sourced from the library of Keio collection (Baba et al., 2006) and strain AL441 was provided by the Imlay lab (Li and Imlay, 2018). The Δ*mscL*::Kan deletion mutation was constructed by following the procedures of recombineering (Datsenko and Wanner, 2000).

**Table S2**: Plasmids employed in this study.

| Plasmids | Description |
|---|---|
| pHYD5001 | A derivative of the plasmid pTrc99 bearing a replacement of the NcoI site to the NdeI site in the multiple cloning site (MCS). The second NdeI site in the plasmid backbone was mutated (Sharma et al., 2016) |
| pRA27 | *alx* cloned in pHYD5001 |
| pMU2385 | Promoter-less plasmid for construction of *lacZ* transcriptional reporter (Praszkier et al., 1992) |
| pMU2386 | Promoter-less plasmid for construction of *lacZ* translational reporter (Lawley and Pitiard, 1994) |
| pRA40 | pMU2385 containing transcriptional reporter of *alx* cloned in the PstI and KpnI sites |
| pRA41 | A derivative of pRA40 bearing a deletion of PRE |
| pRA46 | pHluorin cloned in pHYD5001 |
| pRA48 | pMU2385 containing transcriptional reporter of *mntP* cloned in the PstI and KpnI sites |
| pRA49 | A derivative of pRA40 bearing *alx'*$_{\Delta 16\text{-}53}$ |
| pRA50 | A DNA sequence attached to 5′ of *alx* ORF cloned in the NdeI and HindIII sites of pHYD50001 encoding Alx with HA tag on N-terminus |
| pRA54 | pMU2386 containing translational reporter of *alx* clone in the PstI and BamHI sites |
| pRA55 | A derivative of pRA54 bearing a deletion of PRE |
| pRA56 | A derivative of pRA54 bearing *alx'*$_{\Delta 16\text{-}98}$ |
| pRA57 | pMU2386 containing translational reporter of *mntP* clone in the PstI and SalI sites |
| pRA58 | A derivative pRA50 encoding Alx bearing a D284N mutation |
| pRA61 | A derivative pRA50 encoding Alx bearing a D24N mutation |

| | |
|---|---|
| pRA62 | A derivative pRA50 encoding Alx bearing replacement of D92N |
| pRA63 | A derivative pRA50 encoding Alx bearing replacement of E213Q |
| pRA64 | A derivative pRA50 encoding Alx bearing replacement of D216N |
| pRA65 | A derivative pRA50 encoding Alx bearing replacement of D92C |
| pRA66 | A derivative pRA50 encoding Alx bearing replacement of D216C |
| pRA67 | A derivative pRA50 encoding Alx bearing replacement of D284C |
| pRA69 | A derivative pRA50 encoding Alx bearing replacement of E213C |
| pRA70 | A DNA sequence appended at 5′ of *mntP* ORF cloned in the NdeI and HindIII sites of pHYD5001 encoding HA-tagged version of MntP |
| pRA72 | pHYD5001 containing *mscL* with a DNA sequence abutted at 3′ end cloned in the Nde1 and HindIII sites encoding HA-tagged version of MscL |
| pRA74 | A derivative pRA50 encoding Alx bearing a D73N mutation |
| pRA75 | A derivative pRA50 encoding Alx bearing a E86Q mutation |
| pRA76 | A derivative pRA50 encoding Alx bearing a D222N mutation |

**Table S3**: Oligonucleotide primers employed in this study.

| Primer Designation | Sequence of Primer (5' to 3') | Construction/cloning of |
|---|---|---|
| RAV5 | AAATGCAGGACTGCAGTCTCACCCAGCCGCAGCATATTAAT | pRA40, pRA54 |
| RAV48 | CTTACACATAGGTACCTTAAGCGACAACAACAGCGAATCCGCCCCATAGCAA | pRA40 |
| RAV117 | CTTACACATAGGTACCTTAGCCGACAGTATTCATAG | pRA49 |
| RAV23 | CATGGCATGTGGATCCGCCCCACGACGCCCCTGCAACAACA | pRA54 |
| RAV118 | TGGCATGTGGATCCCCGACAGTATTCATAGAAGTTCC | pRA56 |
| RAV15 | GAATGGTCTGTCATACTTTGTTACCTCCGACGTTGGCCGTTTTTTTATG | pRA41, pRA55 |
| RAV16 | CATAAAAAAACGGCCAACGTCGGAGGTAACAAAGTATGACAGACCATTC | pRA41, pRA55 |
| RAV119 | TAGCAGGACTGCAGAATCACGGACCTGGTACTG | pRA48, pRA57 |
| RAV120 | CTTACACATAGGTACCTTAATCCATCGACATACCAAAC | pRA48 |
| RAV126 | CTCGTTCGGTCGACTCCATCGACATACCAAAC | pRA57 |
| RAV11 | AGGAACTTCTCATATGAATACTGTCGGCACGCCGTTGCTAT | pRA27 |
| RAV12 | GATTAAAAAGCTTTTATCCACCCCGCTGCTTATCATGCCGAT | pRA27, pRA50 |
| RAV139 | GAATTCCATATGTACCCATACGATGTTCCTGACTATGCGGGCGGCCCAAATACTGTCGGCACG | pRA50 |

| | | |
|---|---|---|
| RAV178 | GAATTCCATATGTACCCATACGATGTTCCTGACTATGCGG GCGGCCCAAATATCACTGCTACTGT | pRA70 |
| RAV14 | GGACATTGTCCATATGAATATCACTGCTACTGTTCTTCTT | pRA70 |
| RAV179 | GGAGAATAACATATGAGCATTATTAAAGA | pRA72 |
| RAV180 | GAACATCGTATGGGTATGGGCCAGAGCGGTTATTCTGCTC | pRA72 |
| RAV181 | GATTAAAAAGCTTTACGCATAGTCAGGAACATCGTATGG GTATGGG | pRA72 |
| RAV65 | GCGGATAACAATTTCACACAGGAAACAGCATATGAGTAA AGGAGAAGAACTTTTC | pRA46 |
| RAV66 | AAAATCTTCTCTCATCCGCCAAAACAGCCAAGCTTTTATT TGTATAGTTCATCCATGC | pRA46 |
| RAV144 | CATTATGCTGGCTATCAACCTGTTGTTGCAGGG | pRA61 |
| RAV145 | CCCTGCAACAACAGGTTGATAGCCAGCATAATG | pRA61 |
| RAV146 | AAATCGCTGGCGGTCAATAACGTCTTTGTCTGG | pRA62 |
| RAV147 | CCAGACAAAGACGTTATTGACCGCCAGCGATT | pRA62 |
| RAV148 | GGTACTGATTCTGGTGCAATTGAGCGACGTGATT | pRA63 |
| RAV149 | AATCACGTCGCTCAATTGCACCAGAATCAGTACC | pRA63 |
| RAV150 | TCTGGTGGAATTGAGCAACGTGATTTTCGCCGTGG | pRA64 |
| RAV151 | CCACGGCGAAAATCACGTTGCTCAATTCCACCAGA | pRA64 |

| | | |
|---|---|---|
| RAV152 | CAAGATGCTGATTGTCAACTTCTACCATATTCC | pRA58 |
| RAV153 | GGAATATGGTAGAAGTTGACAATCAGCATCTTG | pRA58 |
| RAV188 | TCGCGCCGTTGCCAATCCACAGGCAC | pRA74 |
| RAV189 | GTGCCTGTGGATTGGCAACGGCGCGA | pRA74 |
| RAV190 | CAGGTTATCTGATTCAGAAATCGCTGGCGG | pRA75 |
| RAV191 | CCGCCAGCGATTTCTGAATCAGATAACCTG | pRA75 |
| RAV194 | GATTTTCGCCGTGAATAGCATTCCG | pRA76 |
| RAV195 | CGGAATGCTATTCACGGCGAAAATC | pRA76 |
| RAV167 | AAATCGCTGGCGGTCTGCAACGTCTTTGTCTGG | pRA65 |
| RAV168 | CCAGACAAAGACGTTGCAGACCGCCAGCGATT | pRA65 |
| RAV169 | GGTACTGATTCTGGTGTGCTTGAGCGACGTGATT | pRA69 |
| RAV170 | AATCACGTCGCTCAAGCACACCAGAATCAGTACC | pRA69 |
| RAV171 | TCTGGTGGAATTGAGCTGCGTGATTTTCGCCGTGG | pRA66 |
| RAV172 | CCACGGCGAAAATCACGCAGCTCAATTCCACCAGA | pRA66 |
| RAV173 | CAAGATGCTGATTGTCTGCTTCTACCATATTCC | pRA67 |

| | | |
|---|---|---|
| RAV174 | GGAATATGGTAGAAGCAGACAATCAGCATCTTG | pRA67 |
| RAV182 | TTAGACTTATGGTTGTCGGCTTCATAGGGAGAATAACATG ATTCCGGGGATCCGTCGACC | MG1655 Δ*mscL*:: Kan |
| RAV183 | TTCTGCTTTCAGGCGCTTGTTAAGAGCGGTTATTCTGCTCT GTAGGCTGGAGCTGCTTCG | MG1655 Δ*mscL*:: Kan |

**Table S4:** Comparison of predicted transmembrane segments (TMS) of Alx using prediction tools[&].

| Tools | TMS 1 | TMS 2 | TMS 3 | TMS 4 | TMS 5 | TMS 6 | TMS 7 | TMS 8 | TMS 9 |
|---|---|---|---|---|---|---|---|---|---|
| DeepTMHMM | 7-27 | 42-62 | 88-101 | 116-136 | 142-160 | 205-228 | 235-255 | 265-283 | 291-311 |
| HMMTOP | 8-27 | 42-64 | 89-108 | 115-137 | 142-161 | 198-221 | 226-245 | 264-283 | 288-307 |
| PHOBIUS | 6-29 | 41-63 | 83-102 | 114-136 | 142-161 | 205-229 | 235-255 | 267-286 | 292-311 |
| TOPCONS | 7-27 | 41-61 | 85-105 | 114-134 | 140-160 | 198-218 | 235-255 | 265-285 | 289-309 |
| OCTOPUS | 7-27 | 41-61 | 85-105 | 113-133 | 135-155 | 198-218 | 236-256 | 265-285 | 287-307 |
| Philius | 8-27 | 42-63 | 84-106 | 114-135 | 142-161 | 199-220 | 235-253 | 264-283 | 290-311 |
| PolyPhobius | 7-29 | 41-63 | 83-106 | 114-136 | 142-160 | 200-228 | 235-255 | 265-283 | 289-311 |
| SCAMP1 | 7-27 | 43-63 | 87-107 | 114-134 | 140-160 | 198-218 | 235-255 | 267-287 | 290-310 |

| | | | | | | | | | |
|---|---|---|---|---|---|---|---|---|---|
| SPOCTOPUS | 7-27 | 41-61 | 85-105 | 114-134 | 136-156 | 198-218 | 236-256 | 265-285 | 287-307 |
| AlphaFold | 7-29 | 37-67 | 69-103 | 108-139 | 143-160 | 204-228 | 234-256 | 265-281 | 289-316 |

[&] DeepTMHMM, HMMTOP, PHOBIUS, TOPCONS, OCTOPUS, Philius, PolyPhobius, SCAMP1, SPOCTOPUS and AlphaFold are described in the refences (Bernsel et al., 2008; Hallgren et al., n.d.; Jumper et al., 2021; Käll et al., 2004; Reynolds et al., 2008; Tsirigos et al., 2015; Viklund et al., 2008; Viklund and Elofsson, 2008)

### Figure S1

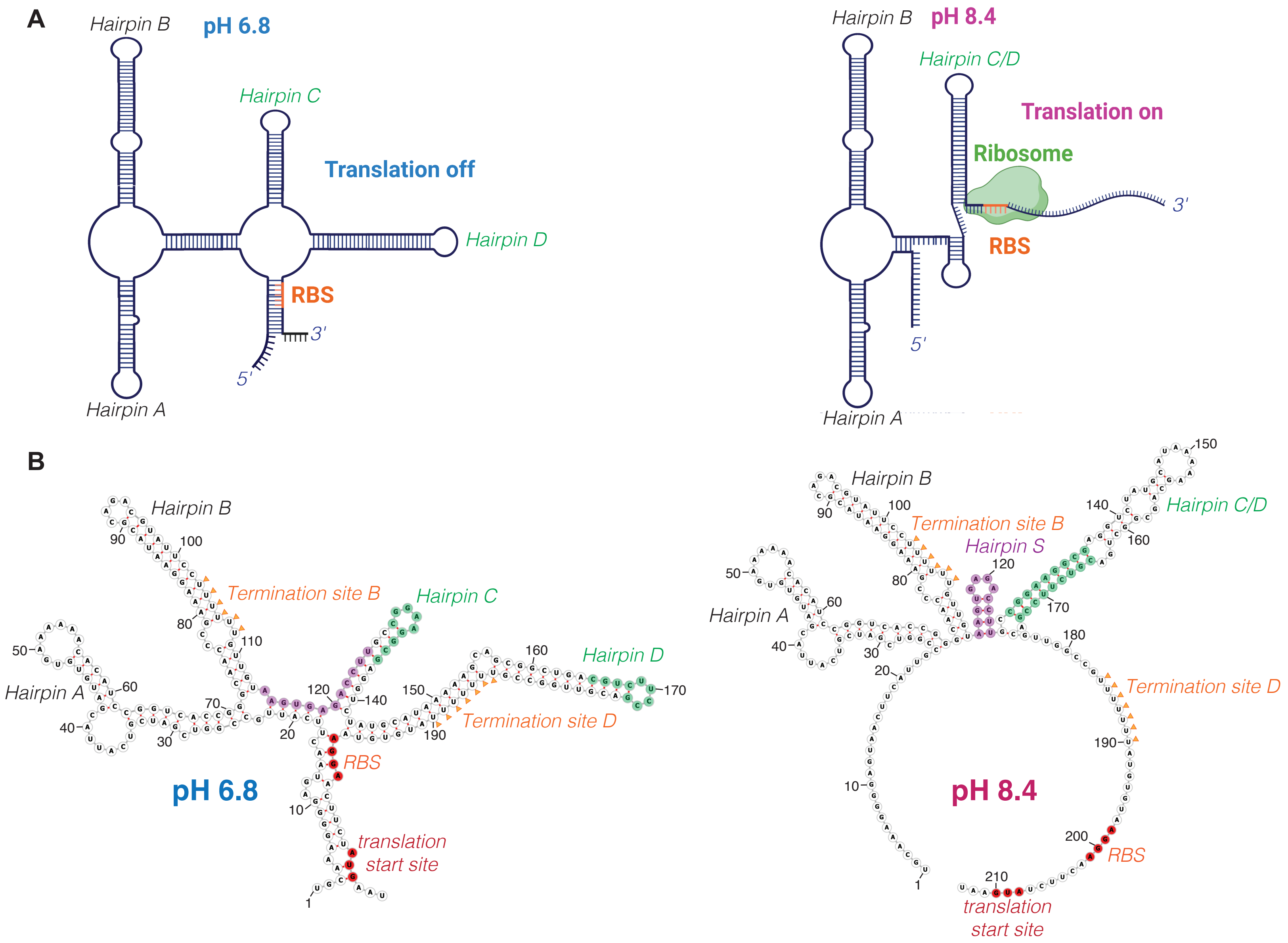


**Figure S1.** Two proposed alternate conformations of PRE-*alx* mRNA. (A) Model proposed by Nechooshtan et al for the two folding outcomes of PRE-*alx* mRNA at different pH. Secondary structure models are based on reporter expression data and RNA structure probing with dimethyl sulfate (DMS) followed by primer extension in two pH conditions. Salient features of the proposal were that during synthesis of PRE-*alx* mRNA by RNA polymerase in alkaline pH, PRE adopts a translationally active structure so that the RBS (in red) becomes available for ribosome to initiate translation, thus turning on the expression of downstream gene (*alx*). However, PRE synthesis results in a translationally inactive structure in neutral pH with the RBS occluded, turning off the expression of *alx* gene. (B) The specific structural elements of PRE-*alx* mRNA involve hairpin C and hairpin D. Alkaline pH favors alternate pairing of upper portion of hairpin C resulting in formation of a putative hairpin S (purple), and lower portion of hairpin C pairs with hairpin D producing C/D hairpin (green).

### Figure S2

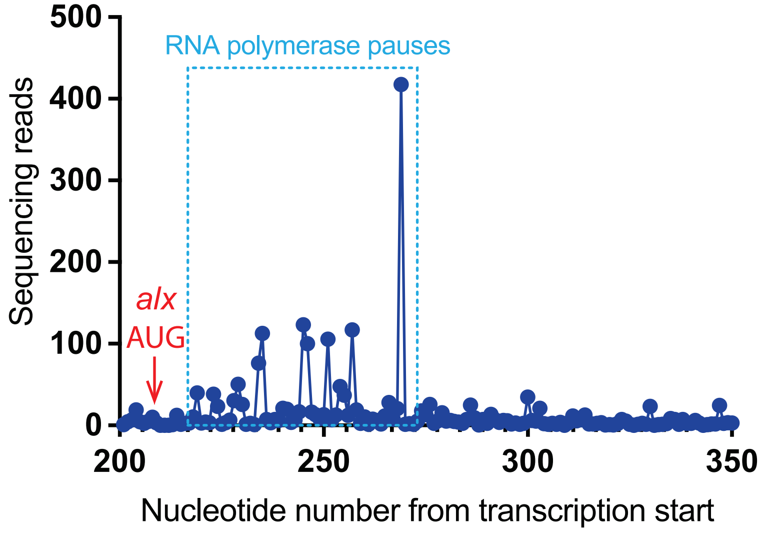


**Figure S2.** The reads for cellular nascent elongating transcripts (NET-seq) from immunoprecipitated RNA polymerases were aligned with respect to transcription start site of *alx*. The translation start site of *alx* and prominent putative transcription pause sites are indicated in the plot.

### Figure S3

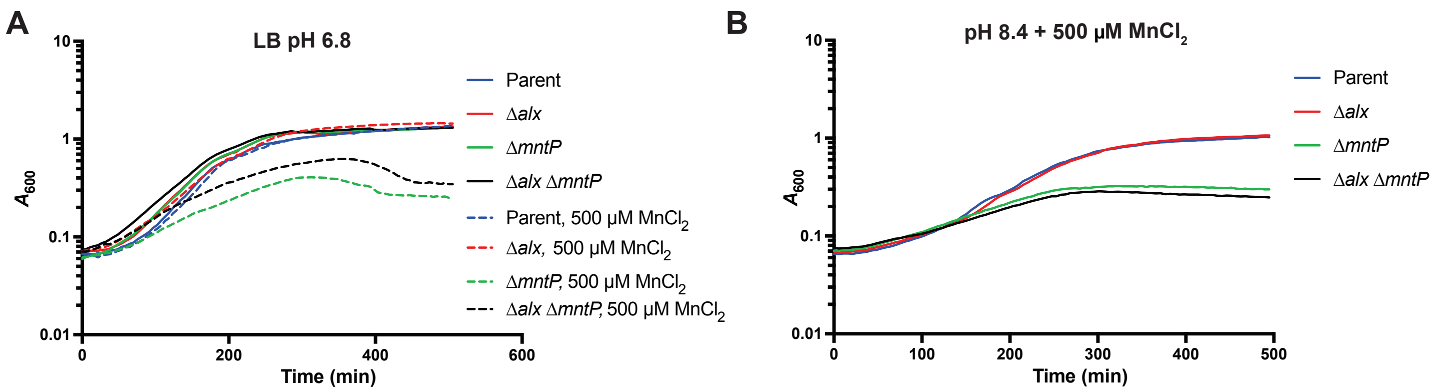


**Figure S3.** Growth of the parent *E. coli* strain (MC4100), its Δ*alx*::Kan (Δ*alx* RAS31), Δ*mntP*::Kan (Δ*mntP* RAS31) and Δ*alx mntP*::Kan (Δ*alx* Δ*mntP* RAS42) derivatives in LB media (pH 6.8) and in LB media (pH 6.8) supplemented with 500 μM $MnCl_2$ (A) and in LBK media (pH 8.4) with supplemented with 500 μM $MnCl_2$ (B).

### Figure S4

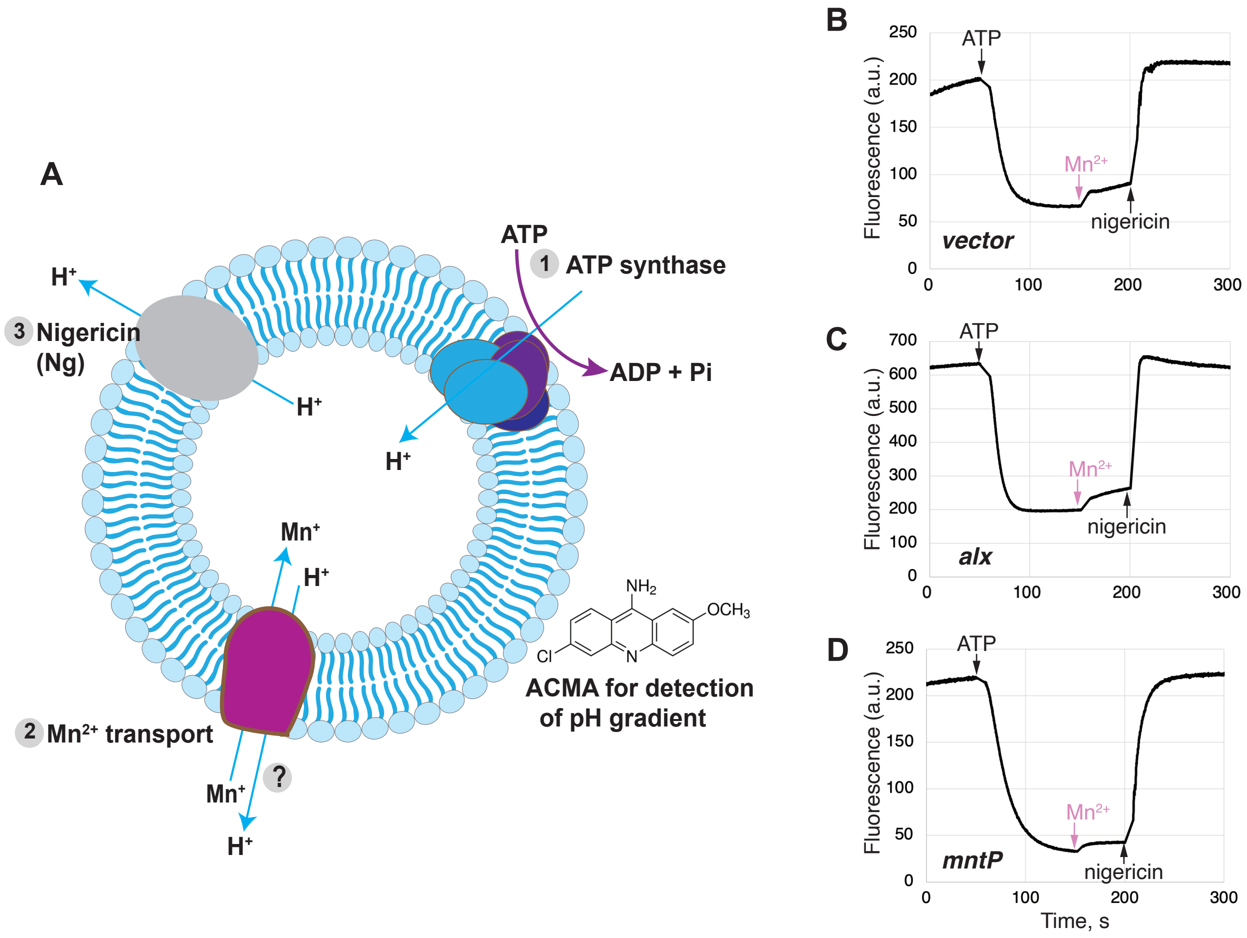


**Figure S4.** Testing for $Mn^{2+}$-induced Alx-mediated proton flux in inside-out vesicles. (A) An overview of experiment. Inside-out vesicles were prepared from RAS42 strain chromosomally defective for *alx* and *mntP* bearing a vector (pHYD5001) or its derivatives expressing $Alx^{HA}$ (pRA54) or $MntP^{HA}$ (pRA70) from $P_{trc}$ promoter. Vesicles were incubated with ACMA, the pH sensitive fluorescent dye. Quenching of ACMA fluorescence was monitored following addition of ATP at 50 s, and then effect of $Mn^{2+}$ addition on recovery of fluorescence was tested at 150 s (panels B, C, and D). Recovery after addition of the uncoupler nigericin (Ng) was recorded at 200 s.

### Figure S5

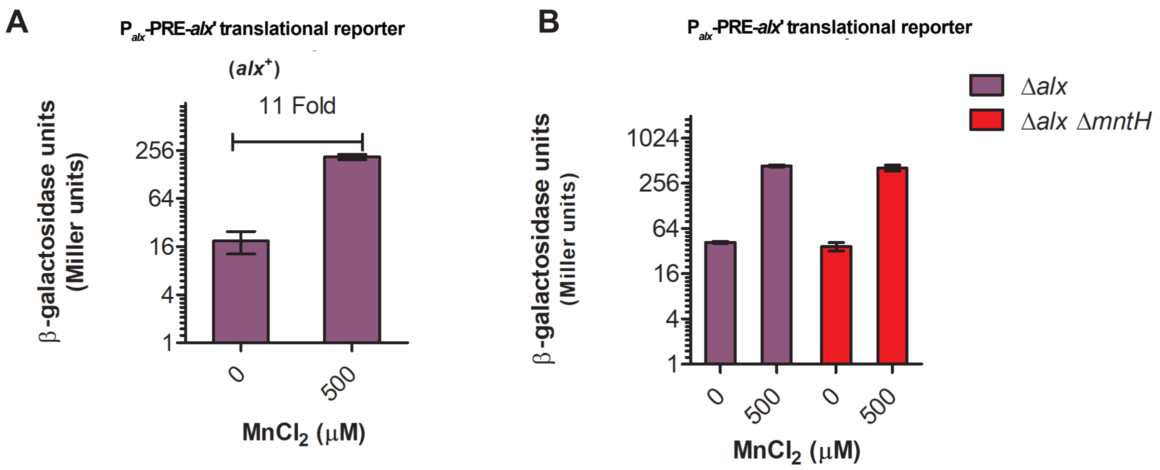


**Figure S5.** β-galactosidase activity (Miller units) as a reporter of *alx* translation ($P_{alx}$-PRE-*alx'*-*lacZ,* pRA54) was measured in mid-log phase grown cultures of *alx*$^{+}$ strain (MC4100) (A) and its Δ*alx*::Kan derivative, RAS40 and Δ*alx mntH*::Kan derivative, RAS93 (B). The cultures were grown in LB media (pH 6.8) with and without supplemented with 500 μM $MnCl_2$. The error shown is standard deviation of three repeats of the experiment.

### Figure S6

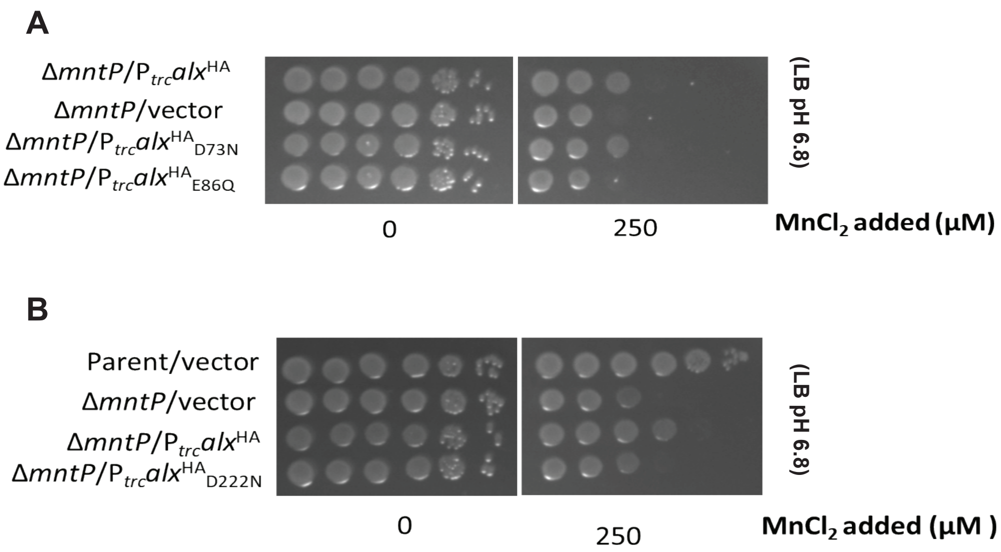


**Figure S6.** (A) Tenfold serial dilutions of overnight grown cultures of Δ*mntP*::Kan mutant (RAS32) bearing a vector (pHYD5001) or a derivative of pHYD5001 expressing $Alx^{HA}$ (pRA50, $Alx^{HA}{}_{D73N}$ (pRA74), and $Alx^{HA}{}_{E86Q}$ (pRA75) from a $P_{trc}$ promoter were spotted on the surface of LB agar (pH 6.8) supplemented with 250 µM $MnCl_2$. (B) Tenfold serial dilutions of overnight grown cultures of parent (MC4100) bearing a vector (pHYD5001) and Δ*mntP*::Kan mutant (RAS32) bearing a vector (pHYD5001) or a derivative of pHYD5001 expressing $Alx^{HA}$ from a $P_{trc}$ promoter (pRA50), $Alx^{HA}{}_{D222N}$ (pRA76) from a $P_{trc}$ promoter were spotted on the surface of LB agar (pH 6.8) supplemented with 250 µM $MnCl_2$.

### Figure S7

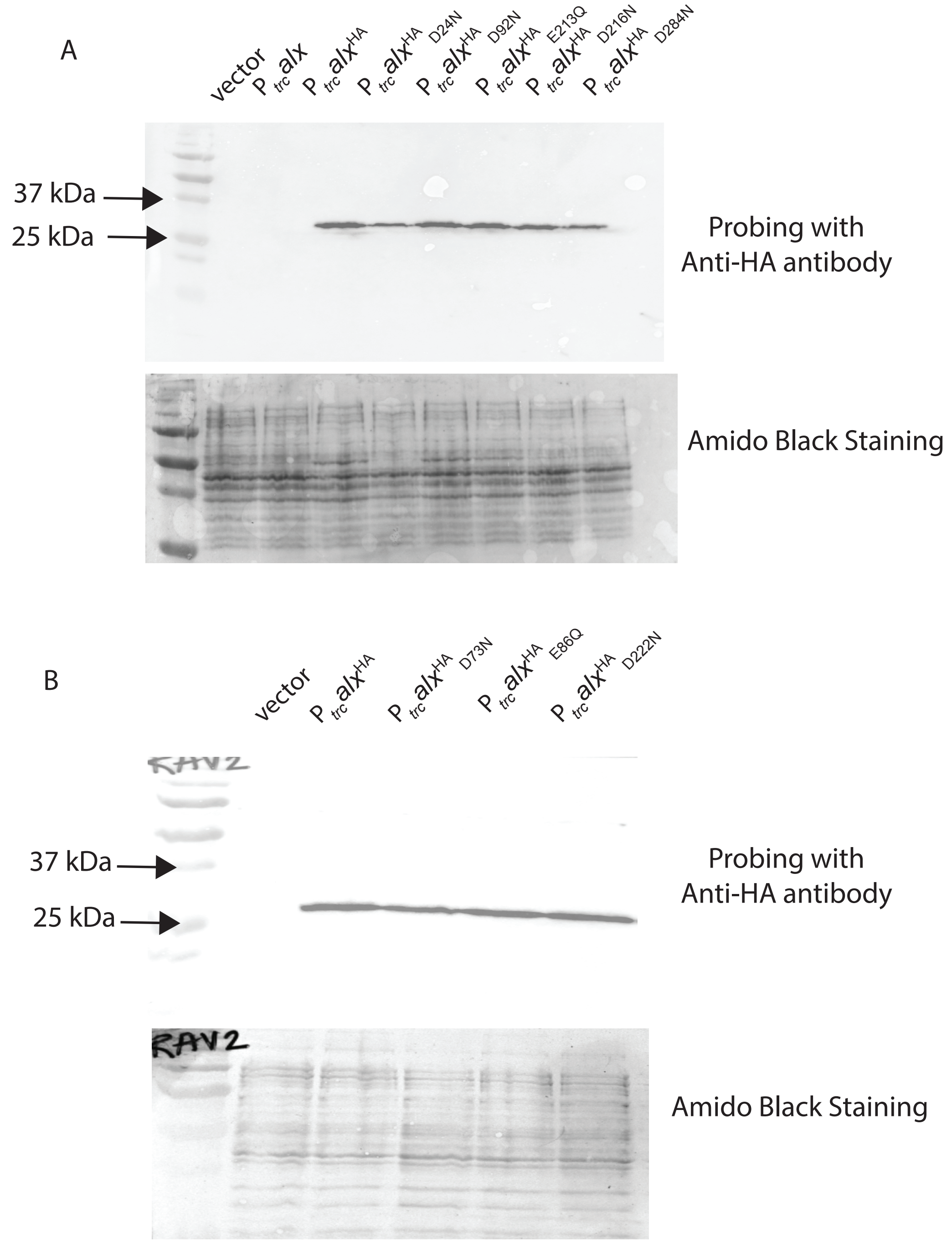


**Figure S7.** (A) Anti-HA immunoblotting to measure the levels of HA-tagged Alx and its variants (D24N, D92N, E213Q, D216N and D284N) expressed from $P_{trc}$ promoter in whole cell extracts of Δ*mntP*::Kan mutant (RAS32). Equal loading of samples across the lanes were gauged with amido black staining of PVDF membrane. (B) Anti-HA immunoblotting to measure the levels of HA-tagged Alx and its variants (D73N, E86Q and D222N) expressed from $P_{trc}$ promoter in whole cell extracts of Δ*mntP*::Kan mutant (RAS32). Equal loading of samples across the lanes were gauged with amido black staining of PVDF membrane.

### Figure S8

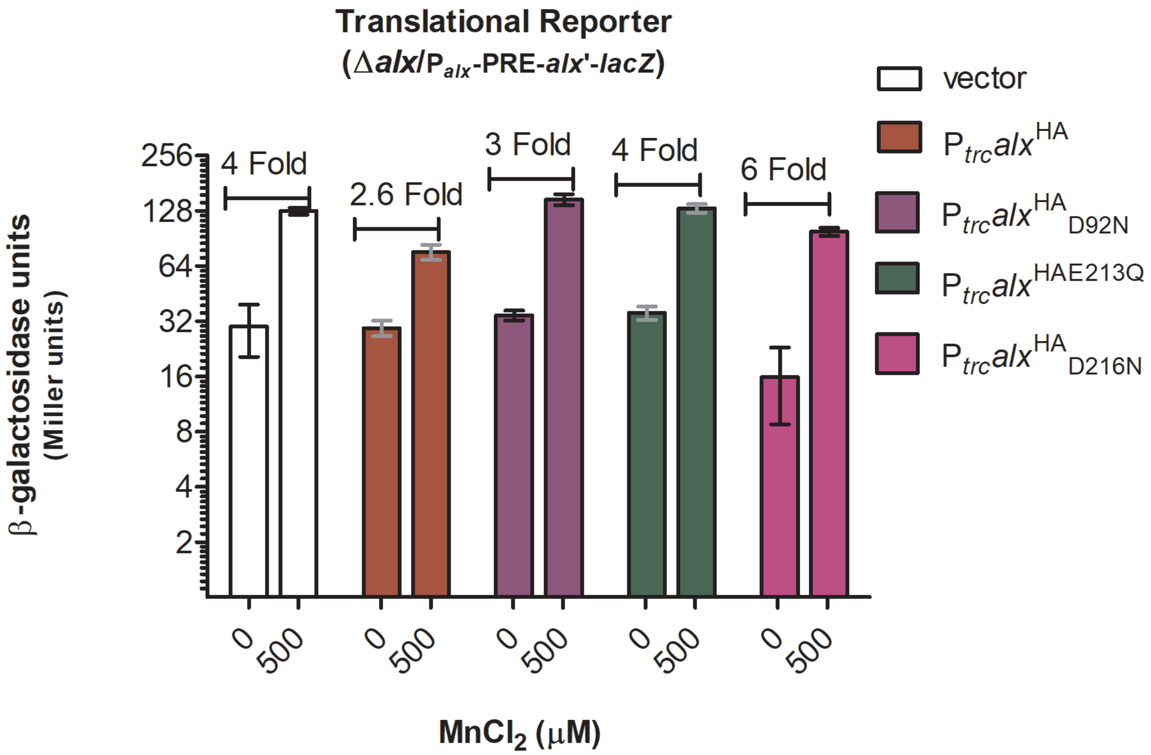


**Figure S8.** β-galactosidase activity (Miller units) as a reporter of *alx* translation ($P_{alx}$-PRE-*alx*'-*lacZ,* pRA54) was measured for mid-log phase grown cultures of Δ*alx*::Kan derivative (RAS31) bearing a vector (pHYD5001) or a derivative of pHYD5001 expressing $Alx^{HA}$ (pRA27), $Alx^{HA}{}_{D92N}$ (pRA62), $Alx^{HA}{}_{E213Q}$ (pRA63), $Alx^{HA}{}_{D216N}$ (pRA64) from $P_{trc}$ promoter. The cultures were grown in LB media (pH 6.8) with and without supplemented with appropriate concentration of ampicillin, $MnCl_2$ and IPTG. The error shown is standard deviation of three repeats of the experiment.

### Figure S9

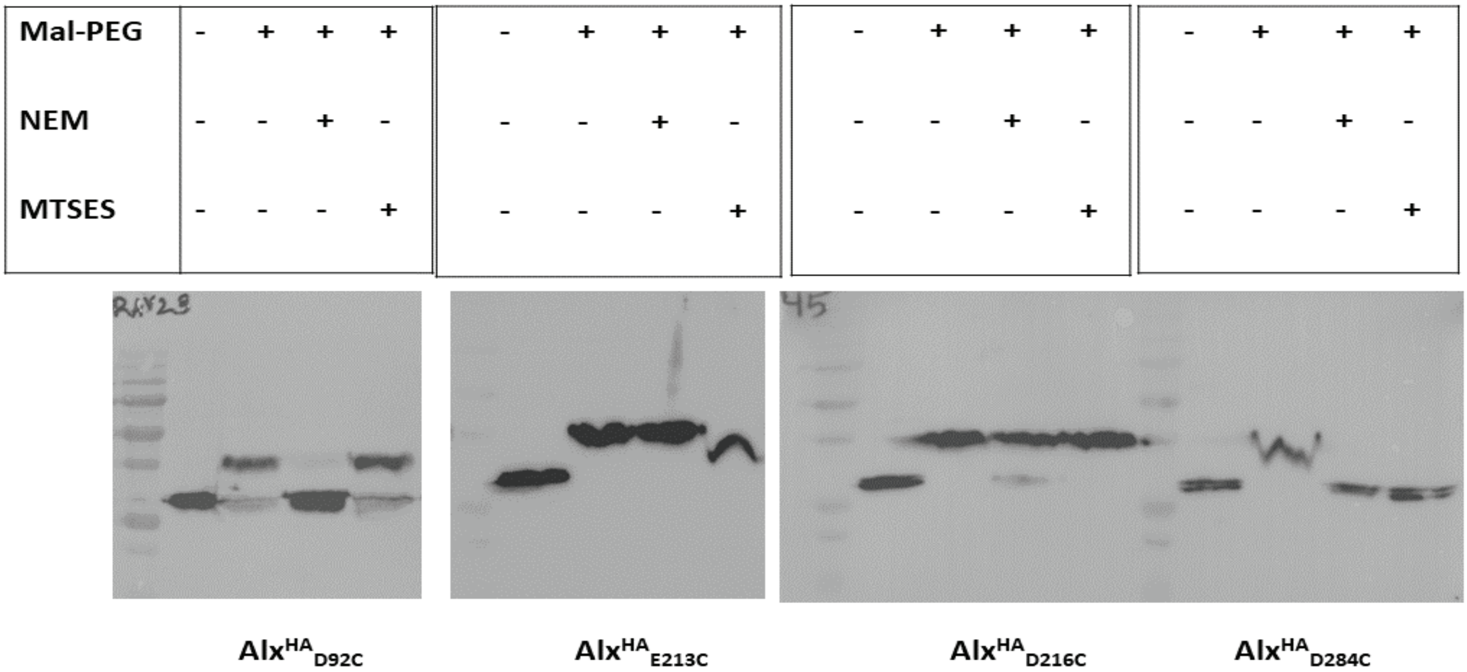


**Figure S9.** Inferring topological locations of important negatively charged residues by substitution with cysteines in Alx. Anti-HA immunoblots to detect the reactivity of mono-cysteine substituted HA-tagged Alx (D92C, E213C, D216C and D284C) with Mal-PEG after an exposure to reagents NEM or MTSES. Samples untreated and treated with only Mal-PEG depicted the effects of NEM or MTSES on cysteine modification based on topological location of the residue.
